# NADP-malic enzyme 1 couples ABA signaling to ROS-auxin patterning to restrict Arabidopsis root growth

**DOI:** 10.64898/2026.03.12.711270

**Authors:** Ying Fu, Maroua Bouzid, Maximilian Klamke, Eva-Maria Schulze Isig, Maximiliano M. Sosa, Gereon Poschmann, Vera Wewer, Sabine Metzger, Mariana Saigo, Veronica G. Maurino

**Affiliations:** Molecular Plant Physiology, Institute of Cellular Molecular Botany (IZMB), University of Bonn, Kirschallee 1, 53115 Bonn, Germany; Centro de Estudios Fotosintéticos y Bioquímicos (CEFOBI-CONICET), University of Rosario, Suipacha 570, 2000 Rosario, Argentina; Molecular Proteomics Laboratory, Biomedical Research Centre (BMFZ) & Institute of Molecular Medicine, Proteome research, Medical Faculty and University Hospital Düsseldorf, Heinrich Heine University Düsseldorf, Universitätsstraße 1, 40225 Düsseldorf, Germany; CEPLAS Plant Metabolism and Metabolomics Facility, Institute for Plant Sciences, University of Cologne, 50674 Cologne, Germany

## Abstract

Abscisic acid (ABA) restricts primary root growth by reshaping reactive oxygen species (ROS) dynamics and hormone signaling at the root apex, yet how cellular reductant supply for redox homeostasis is integrated into this response remains unclear. Here, we show that the cytosolic NADP-dependent malic enzyme 1 (NADP-ME1) is required for full ABA inhibition of Arabidopsis primary root elongation after germination. Three independent *me1* loss-of-function mutants retained significant elongation of primary roots under ABA compared with wild type. In wild type, ABA induced an asymmetric auxin response at the root tip, whereas *me1* roots failed to establish this auxin asymmetry and instead accumulated superoxide, indicating disrupted ROS balance. Pharmacological perturbation of auxin transport and ethylene biosynthesis/signaling attenuated the mutant phenotype, linking NADP-ME1 function to auxin-ethylene interactions during ABA-regulated growth. NADP-ME1 loss amplifies ABA-dependent transcriptional rewiring, including induction of oxidative stress and depression of growth-associated hormone modules. Co-immunoprecipitation coupled with bimolecular fluorescence complementation identified ascorbate peroxidase 1 (APX1) and major latex protein-like 34 (MLP34) as NADP-ME1 interaction partners, suggesting functional coupling between NADPH production and ROS detoxification. Together, our results support a model in which NADP-ME1 shapes the superoxide/H_2_O_2_ balance to control auxin patterning at the root tip and thereby execute ABA-mediated growth inhibition.

## Introduction

Plant root growth reflects a dynamic balance between intrinsic developmental programs and environmental cues (Malamy, 2005; Rellan-Alvarez et al., 2016). During abiotic stresses such as drought and salinity, roots remodel their growth patterns to maintain water and nutrient acquisition (Dinneny, 2019). Central to this coordination is the phytohormone abscisic acid (ABA), a key integrator of developmental and stress signaling. ABA modulates processes from root system architecture and germination to stomatal aperture, aligning plant growth with constraints such as drought, salinity, and nutrient limitation (Hirayama and Shinozaki, 2007; Cutler et al., 2010; Finkelstein, 2013). In the model plant *Arabidopsis thaliana*, ABA signaling is well defined and is largely mediated by the PYR/PYL/RCAR receptor-PP2C-phosphatase-SnRK2 kinase core module. This pathway controls both transcriptional programs and post-translational regulation, thereby tuning growth sensitivity to environmental conditions (Umezawa et al., 2009; Raghavendra et al., 2010; Umezawa et al., 2010).

Root responses to ABA are concentration-dependent: low ABA levels can transiently promote elongation, whereas higher levels inhibit primary root growth (Rodrigues et al., 2009; Li et al., 2017; Sun et al., 2018). At elevated ABA levels, root growth inhibition is associated with reduced cell division in the root apical meristem and suppressed cell expansion in the elongation zone of the roots (Takatsuka and Umeda, 2014). These effects are further modulated by extensive crosstalk with other signaling pathways, including auxin, ethylene, gibberellin (GA), and brassinosteroid (BR), as well as calcium (Ca^2^⁺) and redox signaling (Rodrigues et al., 2009; Thole et al., 2014; Sun et al., 2018; Zluhan-Martínez et al., 2021). In this context, ABA stimulates ethylene biosynthesis via Ca^2^⁺-dependent kinases that phosphorylate and stabilize 1-aminocyclopropane-1-carboxylic acid synthases (Luo et al., 2014). Ethylene then modulates local auxin accumulation, transport, and signaling, with high ABA levels dampening these processes and ultimately leading to growth arrest (Růžička et al., 2007; Thole et al., 2014; Li et al., 2017; Zluhan-Martínez et al., 2021). Crosstalk extends to GA and BR pathways, where ABA antagonizes GA-and BR-dependent elongation by repressing phytochrome interacting factors (PIF) and BR enhanced expression (BEE) transcription factors while activating stress-adaptive genes (de Lucas and Prat, 2014; Gruszka, 2018). Together, these multilayered interactions shape the hormonal landscape that controls root elongation.

A critical component of ABA action is its impact on cellular redox homeostasis. In fact, ABA promotes the generation of reactive oxygen species (ROS) in part through activation of plasma-membrane NADPH oxidases AtRbohD and AtRbohF (Kwak et al., 2003). These enzymes generate superoxide radical (O_2_•⁻), which is rapidly converted into hydrogen peroxide (H_2_O_2_), a key signaling molecule in developmental and stress responses (Hossain et al., 2015). ROS contribute to root growth arrest by decreasing auxin transport, reducing the expression of cell cycle-related genes, altering cellular redox balance, disrupting DNA replication, and damaging cell wall structure (Jiao et al., 2013; Tsukagoshi, 2016; Pasternak et al., 2023). Accordingly, scavenging mechanisms exist to maintain ROS homeostasis in roots and this relies on the activity of antioxidant enzymes such as superoxide dismutase (SOD) and ascorbate peroxidase (APX). SOD converts O_2_•⁻ to H_2_O_2_, and APX detoxifies H_2_O_2_ using ascorbate as an electron donor (Davletova et al., 2005; Hossain et al., 2015). To illustrate how redox homeostasis influences growth, Davletova et al. (2005) showed that mutants lacking cytosolic APX1 exhibit increased oxidative stress and altered root development. Thus, ABA regulates growth not only through modification of gene expression but also by adjusting the O_2_•⁻/H_2_O_2_ ratio that defines cellular signaling thresholds. Changes in redox induced by ABA serve as upstream modulators of hormone activity, establishing a functional bridge between redox state and developmental signaling. In this context, ROS shape auxin distribution by disrupting cytoskeleton-dependent vesicle trafficking of the auxin efflux facilitator PIN in the root apex, thereby influencing asymmetric auxin accumulation (Zwiewka et al., 2019; Pasternak et al., 2023).

At the biochemical level, redox metabolism underpins this regulation via NADPH which serves as the main electron donor that fuels antioxidant systems such as the SOD-APX-glutathione cycle (Davletova et al., 2005; Correa-Aragunde et al., 2013; Hossain et al., 2015 2012; Chen et al., 2019; Sun et al., 2019). NADP-dependent malic enzyme (NADP-ME) catalyzes the oxidative decarboxylation of malate to pyruvate and CO_2_ using NADP⁺ as a cofactor, thereby contributing to cytosolic NADPH homeostasis (Voll et al., 2012; Chen et al., 2019; Sun et al., 2019). In *Arabidopsis* grown under non-stress conditions, NADP-ME1 displays very low basal expression in roots but is strongly induced by multiple stress conditions, similar to its maize ortholog (Detarsio et al., 2008; Arias et al., 2018; Sewelam et al., 2020). Consistently, loss of NADP-ME1 alters tolerance to oxidative and metal stress (Badia et al., 2020) and perturbs ABA-regulated processes, including seed germination (Arias et al., 2018; Yazdanpanah et al., 2019) and the inhibition of primary root growth (Arias et al., 2018).

Despite these links, the mechanistic contribution of NADP-ME1 to ABA-mediated primary root growth inhibition remains unresolved. Here, we set out to define the role of NADP-ME1 within the ABA signaling network. Using a reverse-genetic framework combined with multiple physiological assays, imaging, pharmacological inhibition, and transcriptomic analyses, we examined how NADP-ME1 shapes ROS signatures and hormone interactions that together govern root growth responses to ABA.

## Results

### NADP-ME1 is localized to cytosol

Although Arabidopsis NADP-ME1 is not predicted to contain an organellar targeting signal, its subcellular localization had not been experimentally validated. To address this, we fused the *ME1* coding sequence to GFP at either the N-terminus (pUBQ10::GFP::*ME1*) or C-terminus (pUBQ10::*ME1*::GFP) and expressed the constructs under the control of the UBQ10 promoter in *N. benthamiana* leaf protoplasts. In both fusion configurations, GFP fluorescence was distributed throughout the cytosol, surrounding the nuclei and excluding chloroplasts, as indicated by the lack of overlap with chlorophyll autofluorescence (Fig. 1). A GFP control (pUBQ10::GFP) showed a similar cytosolic pattern, confirming that the localization of NADP-ME1 was not influenced by the position of the fluorescent tag. Together, these data indicate that NADP-ME1 is localized to the cytosol.

**Figure 1.**
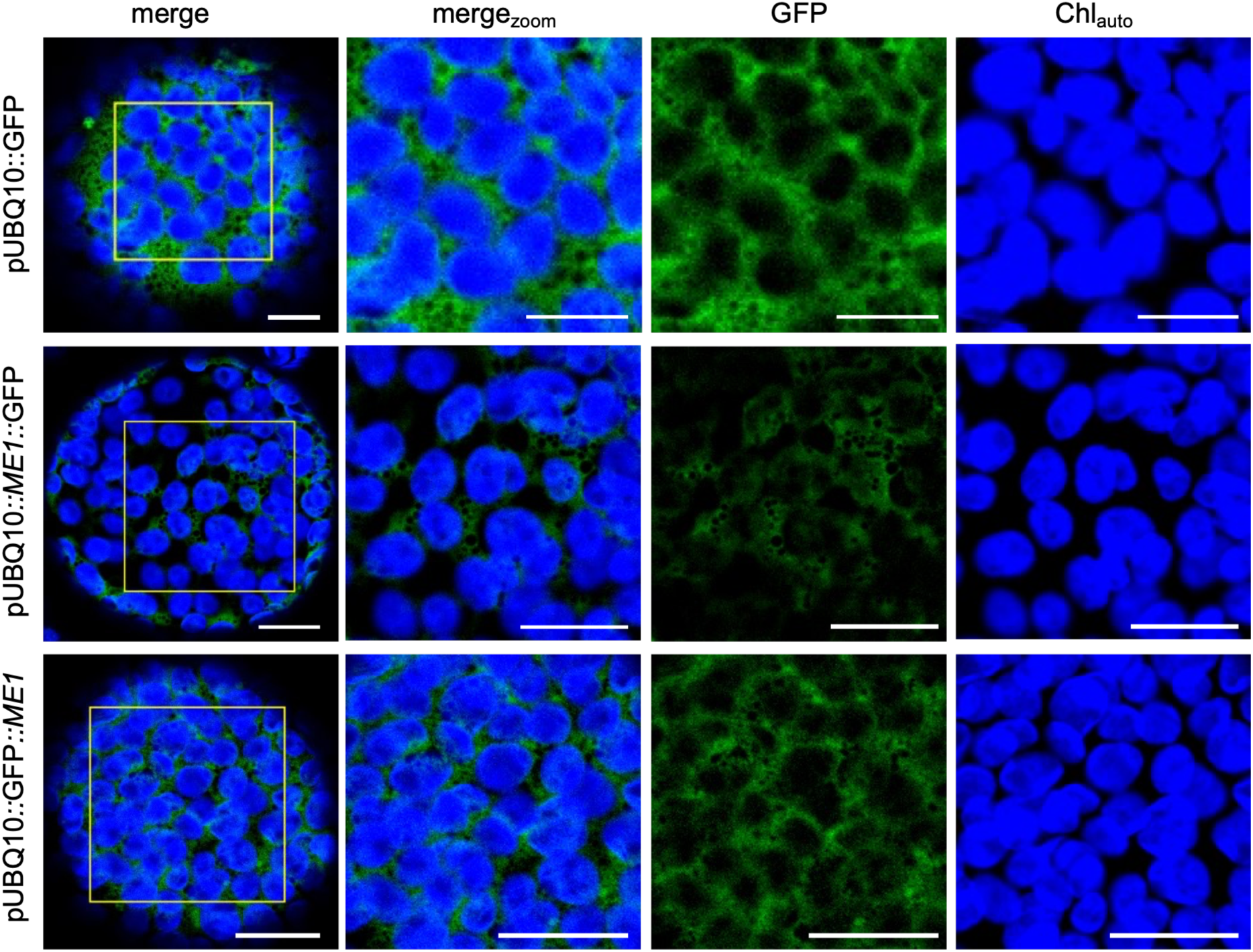
Subcellular localization of *NADP-ME1* in tobacco leaf protoplasts. The coding sequence of *NADP-ME1* was fused in-frame with GFP at either the N-terminus (GFP::*ME1*) or C-terminus (*ME1*::GFP), and expressed under the control of the UBQ10 (AT4G05320) promoter in *Nicotiana benthamiana* leaf protoplasts. GFP alone (pUBQ10::GFP) was used as a control. Images were taken after 3 days of infiltration. merge: overview of the protoplast; merge_zoom_: zoomed overview; GFP: green fluorescence; Chl_auto_: chlorophyll autofluorescence. Scale bars: 10 µm (GFP images); 20 µm (*ME*1::GFP and GFP::*ME1* images).

### Generation and molecular characterization of *NADP-ME1* loss-of-function mutants

Three independent T-DNA insertion lines in the *NADP-ME1* gene, designated *me1.1*, *me1.2*, and *me1.3*, were obtained from the Nottingham Arabidopsis Stock Centre. Genotyping analysis confirmed the presence of T-DNA inserts in the *NADP-ME1* locus in all lines, the precise insertion sites were verified by sequencing, and homozygous plants were selected for further analysis (Suppl. Fig. 1A).

To assess the impact of these insertions on *NADP-ME1* expression, we first examined full-length transcript accumulation in roots by RT-PCR. A clear *NADP-ME1* amplicon was detected in WT roots, whereas the signal was undetectable in all three *me1* mutants (Suppl. Fig. 1B). Quantitative real-time PCR further confirmed the strong reduction of *NADP-ME1* transcript levels in the T-DNA insertion lines (Suppl. Fig. 1C), with all three alleles forming a statistically distinct group with similarly low residual expression. Consistent with these transcriptional defects, an *in-gel* NADP-ME activity assay using seeds imbibed for two days revealed a prominent activity band in WT that was barely detectable in any of the mutant lines (Suppl. Fig. 1D). Together, these data demonstrate that all three mutants are bona fide *NADP-ME1* loss-of-function mutants with effectively abolished NADP-ME1 activity.

### *NADP-ME1* loss-of-function mutants are insensitive to ABA-inhibition of root elongation

Since ABA-responsive promoter motifs in *NADP-ME1* have been identified across different plant species (Tao et al., 2016; Arias et al., 2018), and we previously observed reduced ABA-dependent inhibition of root elongation in a single *me1* mutant line (Arias et al., 2018), we next analyzed the response of independent *me1* alleles to exogenous ABA. We observed similar responses across all three *me1* alleles; therefore, for clarity, we focus on two representative lines in the experiments presented below. Primary root elongation and overall root architecture in the *me1* lines did not differ significantly from WT under control conditions (Fig. 2A). In contrast, after exposure to 1 µM ABA, WT seeds germinated, but further seedling development was strongly inhibited, as indicated by suppressed primary root growth, while *me1* seedlings continued primary root elongation (Fig. 2B). These results indicate that NADP-ME1 is dispensable for root growth under control conditions but is required for proper modulation of ABA-induced inhibition of root elongation during early seedling development.

**Figure 2.**
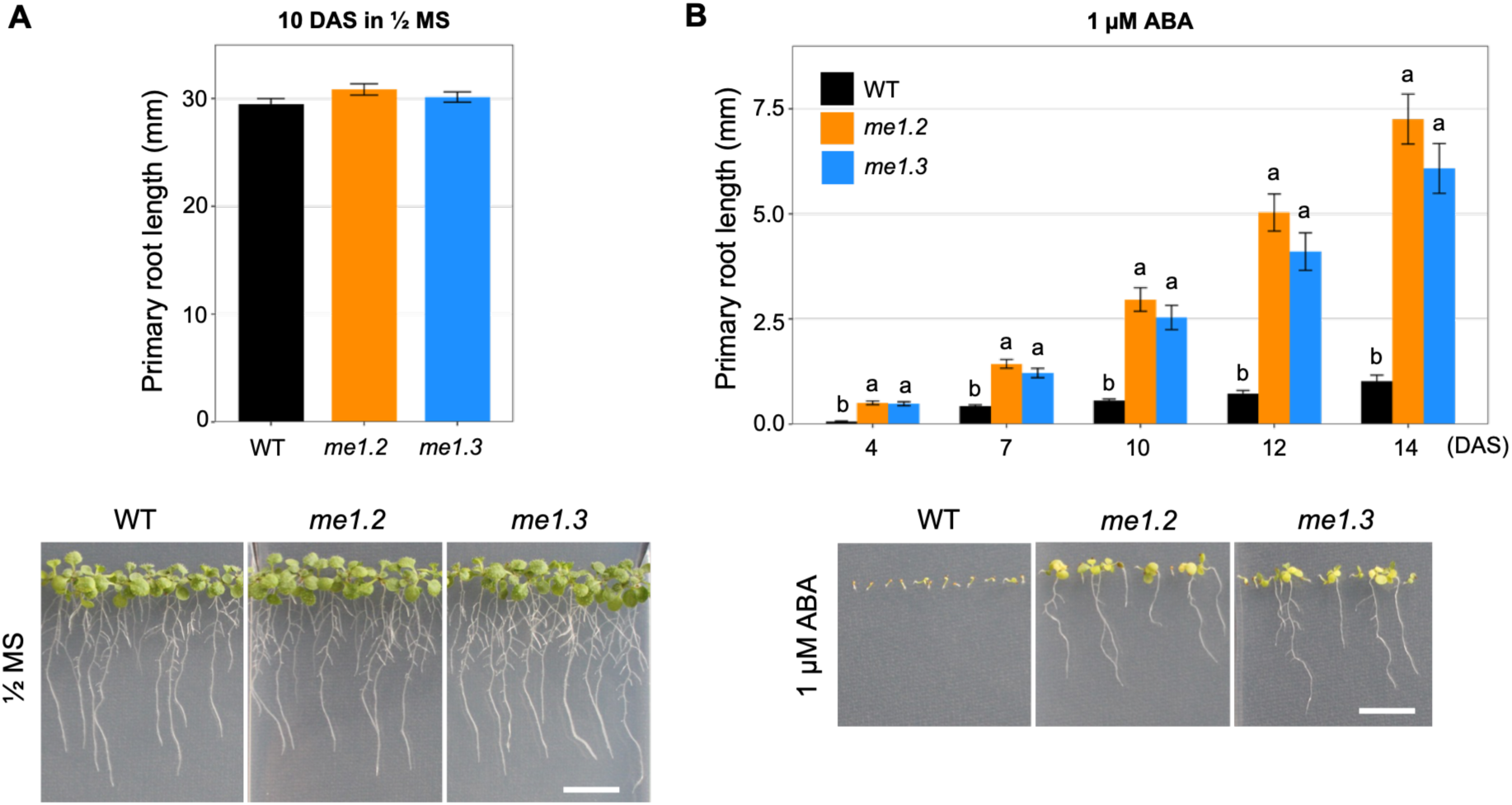
Effect of ABA treatment on root development in *me1* knock-out mutants. **A)** Root growth of WT, *me1.2*, and *me1.3* seedlings on ½ MS medium. Top: Quantification of root length after 10 days of growth. Bottom: Representative images of seedlings grown on ½ MS medium for 10 days (scale bar = 1 cm). Data are presented as mean ± SE (n = 6). **B)** Root growth of WT, *me1.2*, and *me1.3* seedlings on ½ MS medium supplemented with 1 µM ABA. Top: Root length over time (DAS, days after sowing). Bottom: Representative images of seedlings grown on ½ MS medium containing 1 µM ABA for 14 days (scale bar = 1 cm). Data are presented as mean ± SE (n = 6). Root length data were log-transformed prior to analysis. Differences were analyzed using a two-way ANOVA with line and day as factors, followed by pairwise comparisons among lines within each day using estimated marginal means with Sidak adjustment. Different lowercase letters indicate significant differences among lines within the same day (*p* < 0.05).

### Altered auxin distribution in *NADP-ME1* loss-of-function lines under ABA treatment

To test whether NADP-ME1 affects hormone distribution in roots, we introduced auxin and cytokinin reporter constructs into WT and *me1* backgrounds. Auxin responses were monitored using the DR5:RFP reporter. Under control conditions, DR5-driven fluorescence was comparably strong in the root tip and elongation zone of WT and *me1* seedlings, indicating similar auxin response patterns in the absence of stress (Fig. 3A). Upon ABA treatment, DR5:RFP signals in WT became asymmetrically restricted to one side of the root tip, whereas in *me1* lines fluorescence remained symmetrically distributed on both sides of the root apex (Fig. 3A), suggesting an altered ABA-dependent auxin redistribution in the mutants.

**Figure 3.**
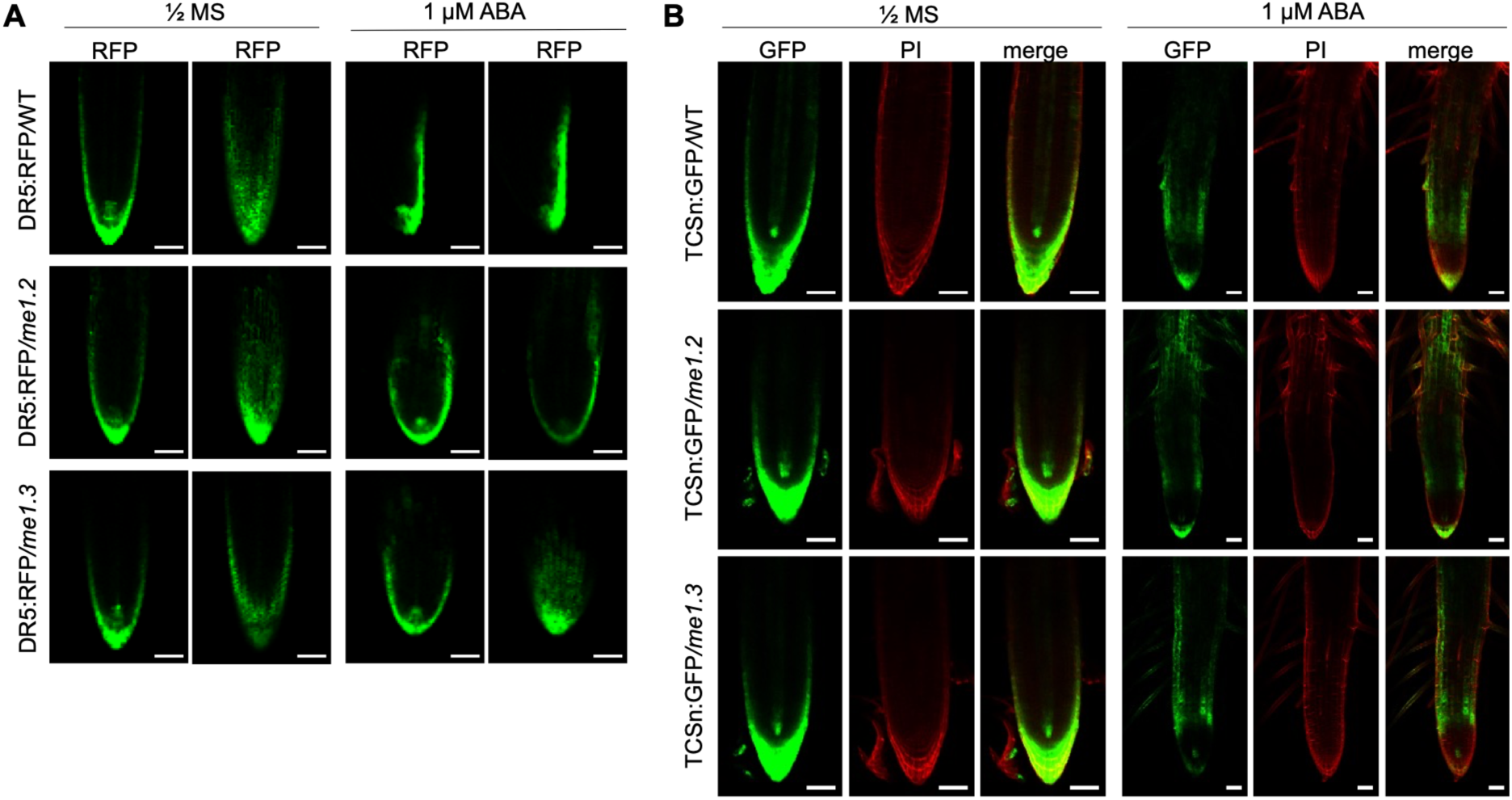
Distribution of auxin and cytokinin in roots of *me1* knockout mutants grown in the absence or presence of 1 µM ABA. **A)** Representative images of DR5:RFP expressed in WT, *me1.2*, and *me1.3* seedlings grown on ½ MS medium without or with 1 µM ABA for 4 days. DR5 is a synthetic auxin-responsive promoter driving RFP expression. RFP, red fluorescence. Scale bar = 50 µm. **B)** Representative images of TCSn:GFP expressed in WT, *me1.2*, and *me1.3* seedlings grown on ½ MS medium without or with 1 µM ABA for 4 days. TCSn (Two Component Signaling - new) is a synthetic cytokinin-responsive promoter driving GFP expression. PI, propidium iodide staining; merge, overlay of fluorescence channels. Scale bar = 50 µm.

Cytokinin signaling was assessed using the TCSn:GFP reporter. Under control conditions, TCSn-driven fluorescence was predominantly detected in the root tip and stele of both WT and *me1* roots (Fig. 3B). Following ABA treatment, cytokinin-responsive signals were reduced to a similar extent in WT and *me1* seedlings, with no obvious differences in spatial distribution (Fig. 3B).

Together, these observations indicate that NADP-ME1 specifically contributes to ABA-dependent re-patterning of auxin signaling in the root apex, whereas ABA-mediated changes in cytokinin responses appear largely unaffected in *me1* mutants.

### Auxin efflux, ethylene and Ca^2^⁺ signaling contribute to NADP-ME1-dependent ABA regulation of root growth

To assess whether auxin, ethylene, or Ca^2^⁺ signaling pathways contribute to NADP-ME1-dependent ABA responses in roots, we used specific inhibitors of auxin transport, ethylene biosynthesis/signaling, and Ca^2^⁺ channels in WT and *me1* mutants. The selection of inhibitors and their working concentrations was informed by previously published studies that demonstrated their efficacy in Arabidopsis (Luo et al., 2014; Li et al., 2017; Cao et al., 2022; Liang et al., 2024). These established concentrations were used as initial benchmarks, after which we conducted preliminary assessments to determine experimental conditions that yielded reproducible and biologically interpretable responses within our system.

Under control conditions, treatment with either the auxin influx inhibitor CHPAA or the efflux inhibitor TIBA reduced primary root elongation to a similar extent in WT and *me1* seedlings, indicating that inhibition of auxin transport alone affects root growth independently of ABA (Fig. 4A; Suppl. Fig. 2A). In co-treatments with ABA and the auxin influx inhibitor CHPAA, root growth responses in both genotypes were comparable to those observed with ABA alone (strong inhibition of primary root elongation in WT seedlings, whereas *me1* mutants displayed reduced sensitivity to ABA-mediated growth inhibition) (Fig. 4B; Suppl. Fig. 2). By contrast, co-treatment with ABA and the auxin efflux inhibitor TIBA resulted in a significant inhibition of primary root elongation in the *me1* mutants, partially restoring their ABA sensitivity and shifting their phenotype towards that of the WT (Fig. 4B; Suppl. Fig. 2). These results indicate that auxin efflux, rather than influx, plays a predominant role in NADP-ME1-dependent ABA regulation of root growth.

**Figure 4.**
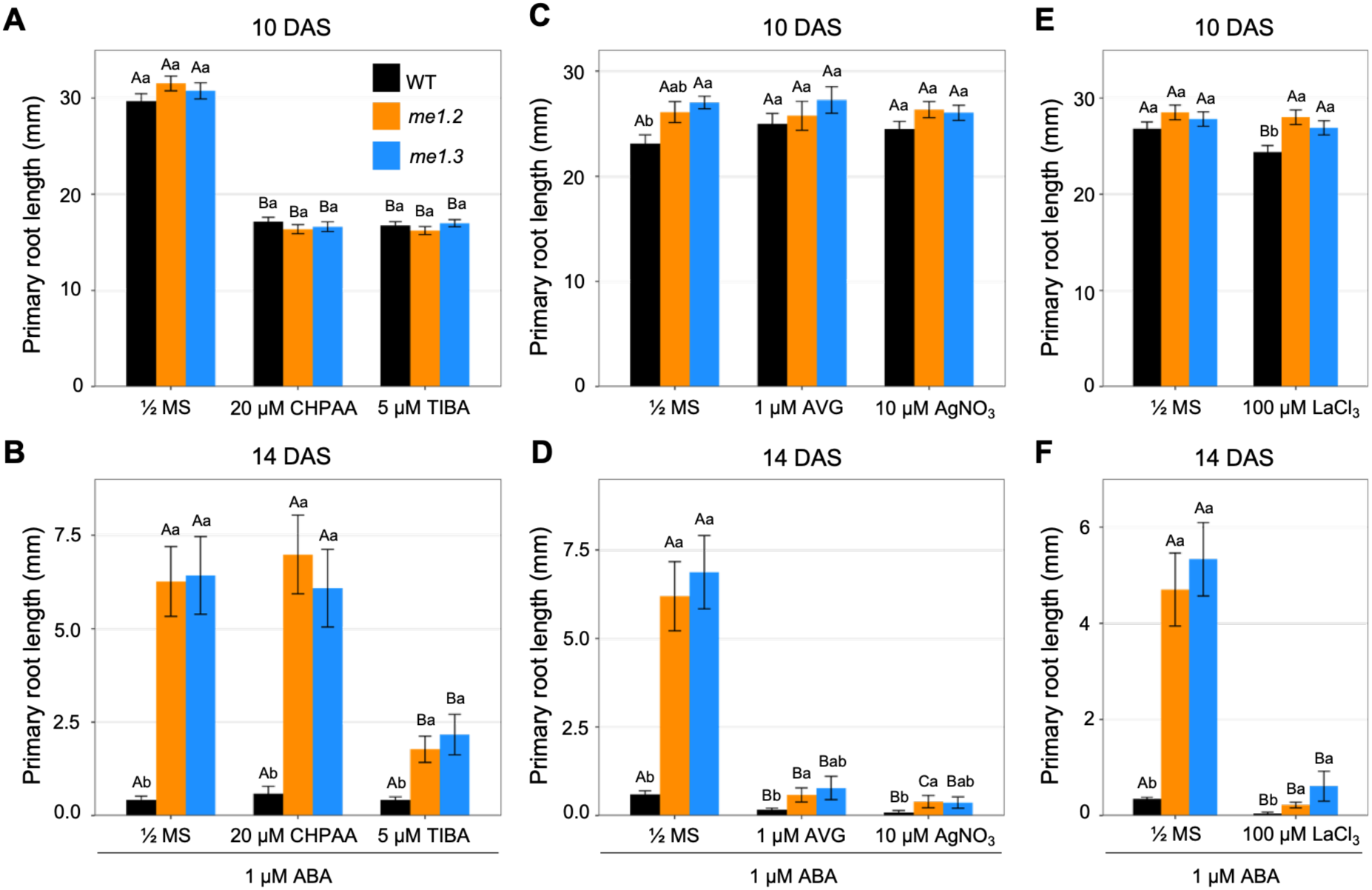
Auxin transport, ethylene, and Ca^2+^ inhibitors differentially affect ABA-dependent root growth in *me1* knock-out mutants. **A)** Primary root length of WT, *me1.2*, and *me1.3* seedlings grown for 10 days on ½ MS medium (control) or ½ MS supplemented with 20 µM CHPAA (3-chloro-4-hydroxyphenylacetic acid; auxin influx inhibitor) or 5µM TIBA (2,3,5-triiodobenzoic acid; auxin efflux inhibitor). **B)** Primary root length of WT, *me1.2*, and *me1.3* seedlings grown for 14 days on ½ MS medium containing 1 µM ABA alone or in combination with 20 µM CHPAA or 5 µM TIBA. **C)** Primary root length of WT, *me1.2*, and *me1.3* seedlings grown for 10 days on ½ MS medium (control) or ½ MS supplemented with 1 µM AVG (L-α-(2-Aminoethoxyvinyl)-glycine hydrochloride; ethylene biosynthesis inhibitor) or 10 µM AgNO_3_ (silver nitrate; ethylene signaling inhibitor). **D)** Primary root length of WT, *me1.2*, and *me1.3* seedlings grown for 14 days on ½ MS medium containing 1 µM ABA alone or in combination with 1 µM AVG or 10 µM AgNO_3_. **E)** Primary root length of WT, *me1.2*, and *me1.3* seedlings grown for 10 days on ½ MS medium (control) or ½ MS supplemented with 100 µM LaCl_3_ (lanthanum chloride; Ca^2+^ channel blocker). **F)** Primary root length of WT, *me1.2*, and *me1.3* seedlings grown for 14 days on ½ MS medium containing 1 µM ABA alone or in combination with 100 µM LaCl_3_. Data are presented as mean ± SE (n = 5). Root length data were log-transformed prior to analysis. Differences were assessed using a two-way ANOVA (line and treatment as factors), followed by pairwise comparisons among lines within each treatment and among treatments within each line using estimated marginal means with Sidak adjustment. Different lowercase letters indicate significant differences among lines within the same treatment, and different uppercase letters indicate significant differences among treatments within the same line (*p* < 0.05).

When ethylene biosynthesis and signaling were blocked by aminoethoxyvinylglycine (AVG) and silver nitrate (AgNO_3_), respectively, primary root growth was not altered under control conditions (Fig. 4C; Suppl. Fig. 3A). In contrast, under co-treatment with ABA and the ethylene inhibitors, root elongation was significantly reduced in all genotypes (Fig. 4D; Suppl. Fig. 3B). These results suggest that NADP-ME1 modulates ABA responses, at least in part, through an ethylene-dependent pathway.

Finally, treatment with the Ca^2^⁺ channel blocker LaCl_3_ did not affect *me1* primary root growth under control conditions (Fig. 4E; Suppl. Fig. 4A). Under co-treatment with ABA and LaCl_3_, however, root elongation was significantly reduced in all genotypes (Fig. 4F; Suppl. Fig. 4B), supporting a contribution of Ca^2^⁺-dependent signaling to ABA-mediated inhibition of root growth.

### Altered auxin and jasmonate homeostasis in *NADP-ME1* loss-of-function mutants

Based on the involvement of auxin and ethylene in the ABA-mediated root growth response, we further quantified endogenous hormone levels in whole seedlings. PCA analysis revealed a clear separation between seedlings grown on ½ MS and those treated with ABA, reflecting an ABA-dependent shift in overall hormone composition (Fig. 5A). The first principal component (PC1, 68.5%) accounted for the major source of variation and clearly separated ABA-treated samples from control. This separation was primarily driven by changes in ABA, including cytokinins (tZ, trans-zeatin; cZ, cis-zeatin; tZR, tZ riboside; tZ7G/tZOG, trans-zeatin-7-glucoside/trans-zeatin-O-glucoside; cZOG/tZ9G, cis-zeatin-O-glucoside/trans-zeatin-9-glucoside; IP, isopentenyladenine; IPR, isopentenyladenosine), the auxin conjugate Ala, indole-3-acetyl-L-alanine (IA), and the jasmonate precursor 12-oxo-phytodienoic acid (OPDA), indicating that ABA treatment strongly alters hormonal homeostasis (Fig. 5A; Suppl. Fig. 5A). The second principal component (PC2, 7.0%) explained a smaller proportion of the variance and distinguished WT from *me1*. This separation was associated mainly with differences in auxin components (IAA, indole-3-acetic acid; IA-Asp, indole-3-acetyl-aspartate; IAM, indole-3-acetamide), cytokinin precursor IPR, and the active jasmonate Jasmonoyl-L-isoleucine (JA-Ile), suggesting that the *me1* mutation affects basal auxin and jasmonate regulation rather than the primary ABA response (Fig. 5A; Suppl. Fig. 5B).

**Figure 5.**
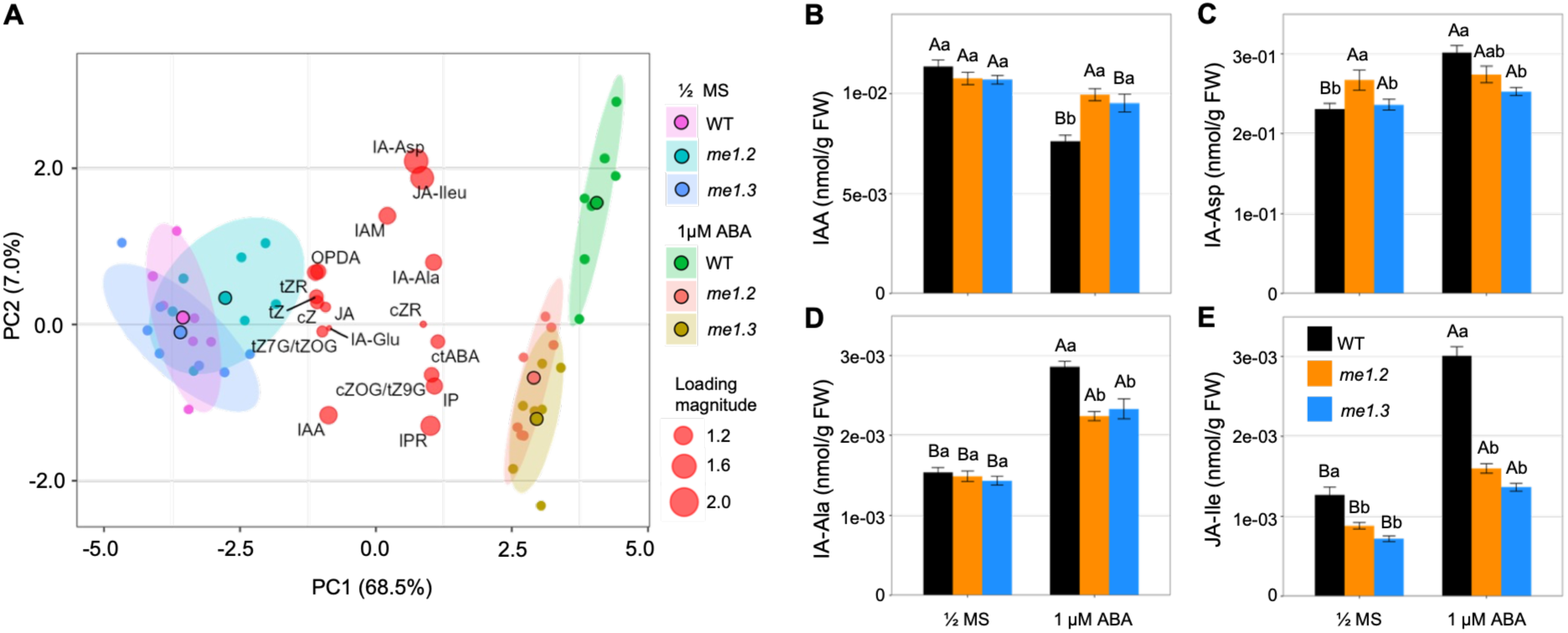
Phytohormone profiles of plants grown in the absence or presence of 1 µM ABA. Seedlings of WT, *me1.2*, and *me1.3* were grown on ½ MS medium without or with 1 µM ABA for two days. Phytohormone content was analyzed using targeted metabolite profiling. **A)** Principal component analysis (PCA) of the phytohormone data illustrating group separation based on genotypes and ABA treatment. Circle size reflects each hormone’s contribution to the principal components, so larger circles represent hormones with greater contributions. Abbreviations: ctABA (cis, trans-ABA); IAA (indole-3-acetic acid); IA-Asp (indole-3-acetyl-aspartate); IA-Ala (indole-3-acetyl-L-alanine); IA-Glu (indole-3-acetyl-glutamate); IAM (indole-3-acetamide); cZ (cis-zeatin); tZ (trans-zeatin); cZR (cZ-riboside); cZR (tZ-riboside); tZ7G/tZOG (trans-zeatin-7-glucoside/trans-zeatin-O-glucoside); cZOG/tZ9G (cis-zeatin-O-glucoside/trans-zeatin-9-glucoside); IP (isopentenyladenine); IPR (isopentenyladenosine); JA (jasmonic acid); JA-Ile (jasmonate Jasmonoyl-L-isoleucine); OPDA (12-oxo-phytodienoic acid). **B)** IAA content. **C)** IA-Asp content. **D)** IA-Ala content. **E)** JA-Ile content. Data represent mean ± SE (n = 7). Differences were assessed using a two-way ANOVA (line and treatment as factors), followed by pairwise comparisons among lines within each treatment and between treatments within each line using estimated marginal means with Sidak adjustment. Different lowercase letters indicate significant differences among lines within the same treatment, and different uppercase letters indicate significant differences among treatments within the same line (*p* < 0.05).

Under ABA treatment, free IAA levels were significantly higher in *me1.2* and *me1.3* than in WT (Fig. 5B), whereas the IAA conjugates IA-Ala and IA-Asp were reduced in both mutants (Fig. 5C, D). By contrast, indole-3-acetyl-glutamate (IA-Glu) levels were slightly, but consistently, higher in the mutants compared with WT (Suppl. Fig. 6). This profile suggests that the elevated free IAA in *me1* alleles arises, at least in part, from altered auxin conjugation dynamics rather than from a general increase in IAA biosynthesis.

For cytokinins, no significant differences among genotypes were detected in the levels of free bases (tZ, cZ) or in the conjugated zeatins *cis*-zeatin riboside (cZR) and tZR, tZ7G/tZOG and cZOG/tZ9G (Suppl. Fig. 6), indicating that ABA-induced cytokinin homeostasis is largely preserved in *me1* backgrounds. Among jasmonates, jasmonic acid (JA) and OPDA levels remained unchanged, whereas JA-Ile, the bioactive form of JA, was significantly lower in *me1.2* and *me1.3* than in WT (Fig. 5E; Suppl. Fig. 6). The reduction in JA-Ile suggests attenuated JA signaling and impaired ABA-JA crosstalk in the mutants, which may contribute to their altered ABA response phenotype. Together, these results indicate that ABA treatment strongly alters hormonal homeostasis, and that the *me1* mutation specifically modulates basal auxin and jasmonate regulation.

### ABA induces ROS accumulation in *NADP-ME1* loss-of-function mutants without altering their oxidative stress sensitivity

To examine whether NADP-ME1 influences redox homeostasis during ABA responses, superoxide accumulation in roots was monitored by histochemical and fluorescent staining. Under control conditions, only low background levels of superoxide (O_2_^•⁻^) were detected in both WT and *me1* root tips (Fig. 6A). By contrast, ABA treatment triggered a marked increase in superoxide accumulation specifically in *me1.2* and *me1.3* root tips, whereas WT showed no detectable change, indicating that NADP-ME1 contributes to the reduced production of ABA-induced O_2_^•⁻^ production (Fig. 6B).

**Figure 6.**
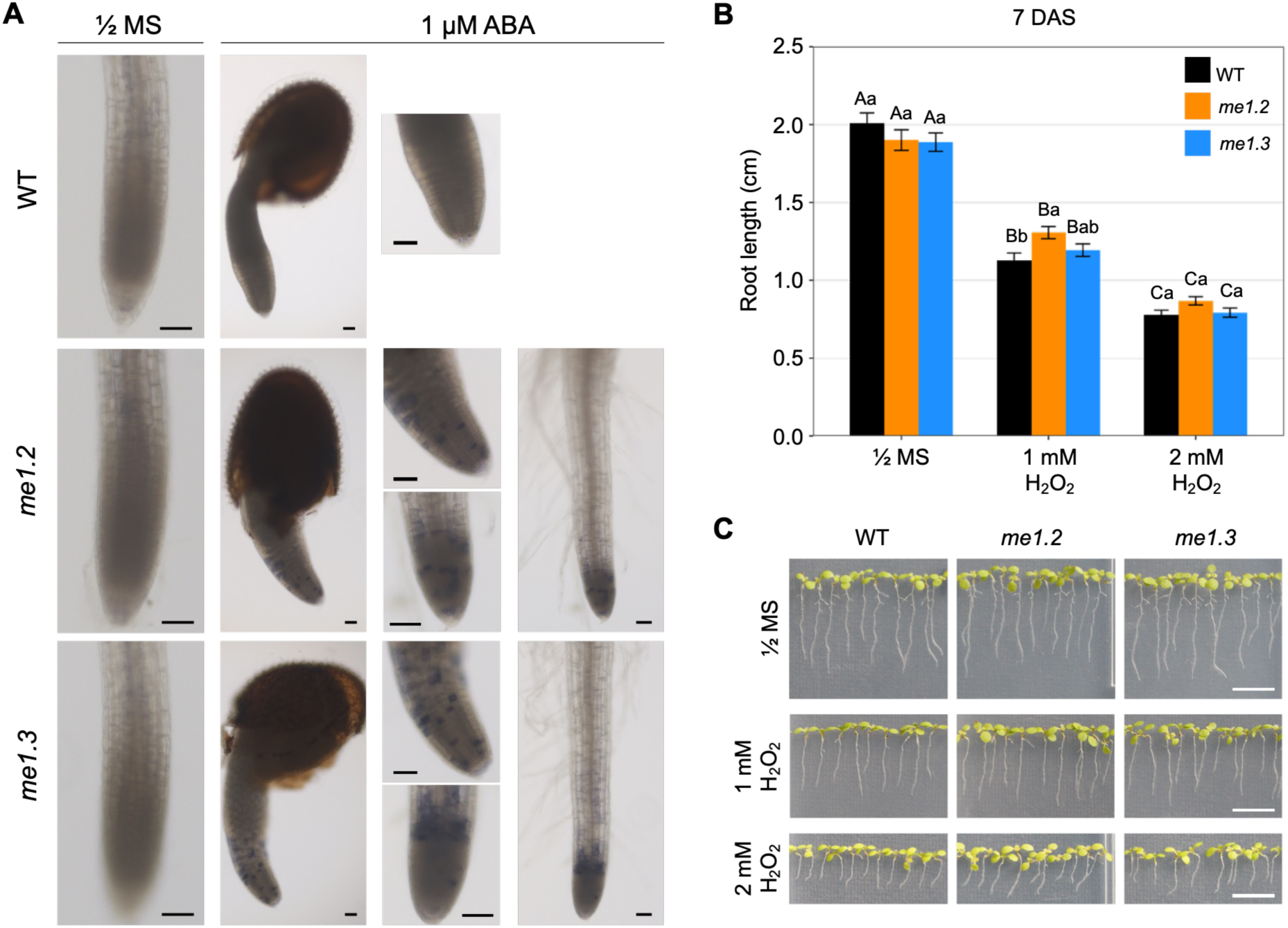
Superoxide accumulation and hydrogen peroxide sensitivity. **A)** Superoxide (O_2_^•⁻^) accumulation detected by nitro blue tetrazolium (NBT) staining in the root tip of four-day-old WT and *me1* seedlings grown on ½ MS medium in the absence or presence of 1 µM ABA. Scale bar: 50 µm. **B)** Effect of H₂O₂ on primary root growth. Root length of WT and *me1* seedlings grown on ½ MS medium (control) or ½ MS supplemented with 1 or 2 mM H_2_O_2_ for 7 days. Data are shown as means ± SE (n = 4). Differences were assessed using a two-way ANOVA (line and treatment as factors), followed by pairwise comparisons among lines within each treatment and among treatments within each line using estimated marginal means with Sidak adjustment. Different lowercase letters indicate significant differences among lines within the same treatment, and different uppercase letters indicate significant differences among treatments within the same line (*p* < 0.05). **C)** Representative images of WT, *me1.2*, and *me1.3* seedlings grown for 7 days on ½ MS medium or ½ MS supplemented with 1 or 2 mM H_2_O_2_. Scale bar: 1 cm.

To assess whether this altered ROS pattern translates into differences in oxidative stress tolerance under control conditions, seedlings were exposed to increasing concentrations of H_2_O_2_. Root elongation declined progressively with higher H_2_O_2_ levels in all genotypes, but no significant differences were observed between WT and *me1* mutants (Fig. 6B, C). These results suggest that loss of NADP-ME1 specifically compromises ROS homeostasis in the context of ABA signaling, without broadly altering sensitivity to exogenous H_2_O_2_.

### Transcriptomic analysis links NADP-ME1 deficiency to altered ABA, redox, and growth signaling

To identify transcriptional changes associated with the ABA-insensitive root growth phenotype of *me1* mutants, we carried out transcriptomic profiling of wild-type and *me1.2* seedlings under control conditions and following ABA treatment. Seedlings were sampled at equivalent developmental stages to ensure that observed transcriptional differences reflected genotype-specific ABA responses rather than variations in developmental progression.

Principal component analysis showed that ABA treatment was the dominant source of variation, with PC1 explaining 97% of the variance and clearly separating ABA-treated samples from controls in both genotypes (Fig. 7A). PC2 accounted for only 1% of the variance and primarily distinguished WT from *me1* within each condition. Across all comparisons, 12,123 genes displayed significant differential expression (|log_2_FC| > 1, padj < 0.05) (Fig. 7B). ABA induced a strong transcriptional response in both genotypes, with 6,049 DEGs in WT (3,902 downregulated, 2,147 upregulated) and 6,074 DEGs in *me1* (3,645 downregulated, 2,429 upregulated) (Fig. 7B). The ABA-responsive gene sets overlapped extensively between genotypes, with similar DEG numbers and comparable fold-change magnitudes (Suppl. Data 1). On the other hand, inter-genotypic comparisons showed that under control conditions, only 91 (17 up, 74 down) genes were differentially expressed between *me1* and WT, suggesting that NADP-ME1 loss has a limited impact on basal gene expression. In contrast, when *me1* and WT seedlings were compared under ABA treatment, 497 genes (216 down, 281 up) were differentially expressed.

**Figure 7.**
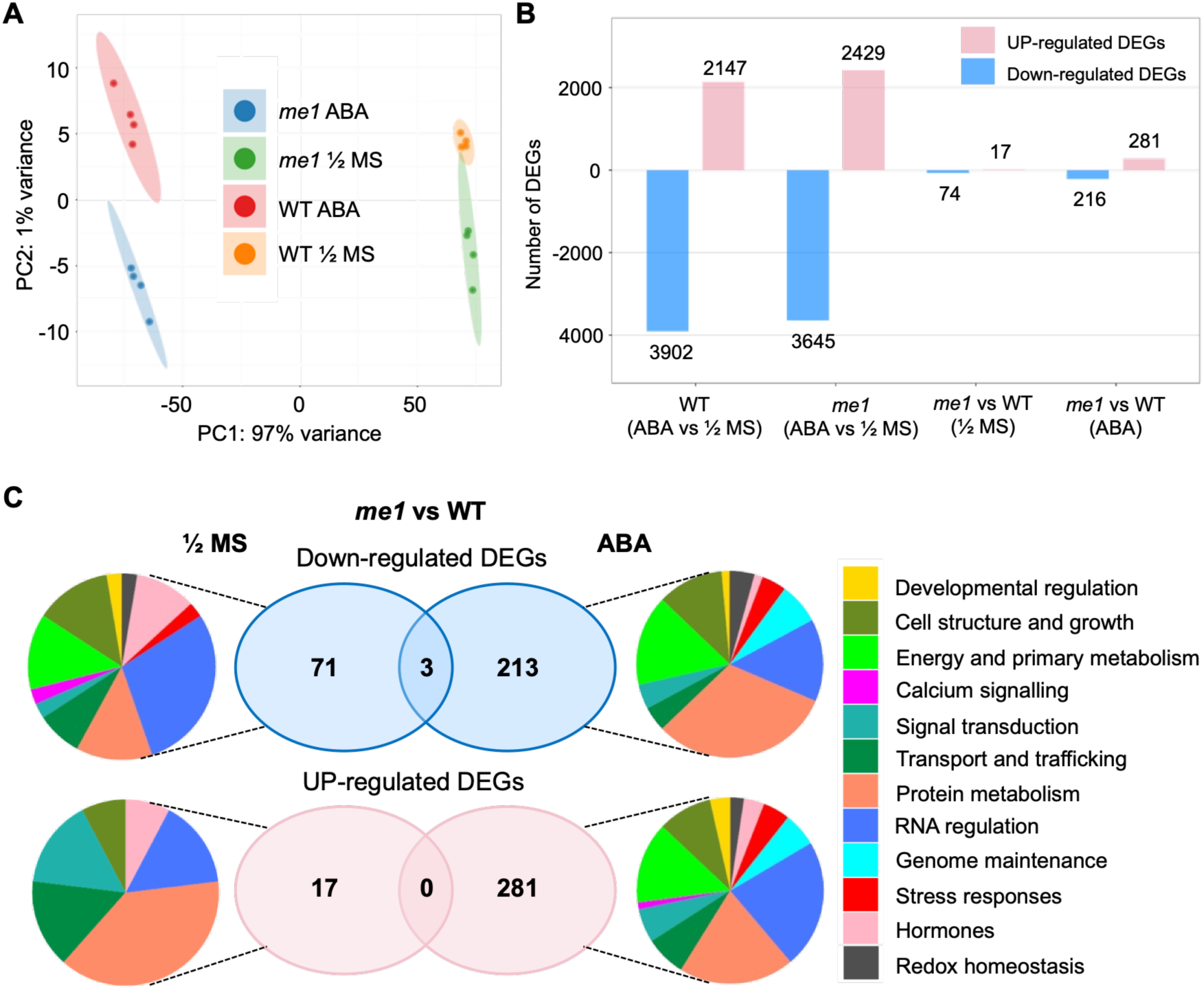
Transcriptomic overview of WT and *me1* seedlings grown in the absence and presence ABA. **A)** Principal component analysis (PCA) of RNA-seq samples based on rlog-transformed read counts, showing the distribution of four biological replicates according to genotypes (WT, *me1*) and treatments (½ MS (control) and ½ MS supplemented with ABA (ABA)). **B)** Bar plot showing the number of significantly up-regulated (pink) and down-regulated (blue) differentially expressed genes (DEGs; (|log_2_FC| > 1, padj < 0.05)) for each comparison (WT ABA vs ½ MS; *me1* ABA vs ½ MS; *me1* vs WT in ½ MS and *me1* vs WT with ABA). **C)** Venn diagrams showing the up-and down-regulated DEGs in *me1* vs. WT in ½ MS (left) and ABA treatment (right), with the corresponding MapMan functional enrichment for each condition displayed as pie charts.

Because genotype-specific ABA-responsive DEGs likely reflect ME1-dependent fine-tuning of ABA signaling, we analyzed genes uniquely regulated between genotypes under control and ABA conditions (Fig. 7C). Among downregulated genes, only 3 of 213 ABA-responsive DEGs overlapped with control-responsive genes, indicating that 98.6% were ABA-specific. Notably, none of the 281 ABA-upregulated genes overlapped with control-upregulated genes, indicating strong ABA-specific regulation.

### MapMan enrichment of genotype-specific DEGs

MapMan enrichment of genotype-specific DEGs indicates that NADP-ME1-dependent transcriptional reprogramming is focused on a discrete set of functional modules (Fig. 7C). In both the ABA_DOWN and ABA_UP gene sets (Fig. 7C), the dominant categories are protein metabolism, energy and primary metabolism, and transcriptional (RNA) regulation. A notable ABA-specific feature is the emergence of the genome maintenance category. In addition, stress response and redox homeostasis are more strongly enriched under ABA than in the corresponding control charts. Within the enriched stress-response genes, we identified plant defensins, germin-like proteins, and cold-acclimation-related members. This category also includes two genes linked to gravitropic signaling: *RG1* (AT1G68370; ∼2.3-fold change; Suppl. Data 2), which encodes a DnaJ-domain protein involved in gravity-signal transduction (Harrison and Masson, 2008), and *LZY1* (AT5G14090; ∼5-fold upregulated, Suppl. Data 2), a regulator of auxin**-**dependent gravitropic responses (Taniguchi et al., 2017). The redox homeostasis module includes genes associated with glutathione and tocopherol metabolism. Strikingly, *FSD1* (AT4G25100), encoding iron superoxide dismutase 1, is downregulated by ∼10-fold in *me1* (Suppl. Data 2). Because FSD1 contributes to superoxide detoxification and supports root development (Dvořák et al., 2021) its repression is consistent with the elevated superoxide signal detected in *me1* root tips under ABA and may contribute to downstream effects on growth-related processes (e.g., redox-sensitive wall remodeling) during stress.

Under control conditions, MapMan enrichment reveals a limited but distinct reorganization of basal transcriptional programs in *me1* compared with WT (Fig. 7C). The most prominent feature of the down-regulated gene set is the dominance of RNA regulation, accompanied by coordinated repression of protein primary metabolism, cell structure and growth and hormone-related functions. Notably, several categories detected only in the down-regulated pie, such as calcium signaling, stress responses, developmental regulation, and redox homeostasis, point to a broad attenuation of regulatory and homeostatic processes under non-stress conditions. Although the hormone category is generic at the bin level, gene-level inspection (Suppl. Data 2) reveals strong repression of specific signaling pathways, including cytokinin signaling via *LOG5* (AT4G35190; ∼36-fold decrease) and jasmonate signaling via *JAZ1* (AT1G19180; ∼5-fold decrease). In contrast, the up-regulated control pie is dominated by protein metabolism, with additional enrichment of RNA regulation and signal transduction. Within the hormone-related category, strong induction of the DELLA protein *RGL2* (AT3G03450; ∼13.5-fold increase) suggests modulation of GA-associated transcriptional regulators in *me1* under basal conditions. Together, these patterns indicate that NADP-ME1 deficiency under control conditions primarily affects regulatory and metabolic balance rather than producing visible developmental alterations.

### Manual curation of highly regulated ABA-responsive DEGs

Because the inter-genotypic DEG sets capture genotype-specific transcriptional responses to ABA, a substantial fraction of DEGs did not fall into statistically enriched MapMan bins (69.8% and 67.1% unassigned among up-and down-regulated genes in *me1* vs WT under ABA, respectively). This outcome is expected for focused and functionally diverse gene sets, for which bin-based overrepresentation tests may lack sensitivity despite robust gene-level regulation (Huang et al., 2009; Khatri et al., 2012). Therefore, we further focused on biological interpretation by manually annotating the most strongly regulated genes in the ABA-treated condition (Suppl. Data 2).

Among the most highly induced genes in *me1* under ABA, the majority map to unassigned MapMan bins yet encode functions consistent with sustained root growth under stress. These include genes implicated in cell wall-mediated growth control, such as a RALF-family signaling peptide (AT3G29780; ∼56-fold increase), previously shown to modulate apoplastic alkalinization and modulate root growth (Abarca et al., 2021), and enzymes associated with carbohydrate and cell wall metabolism, including a GT14-family glycosyltransferase (AT4G03340; ∼52-fold increase) and a cell-wall/apoplast-localized nucleoside phosphorylase (AT4G24350; ∼56-fold increase). In parallel, ABA treatment induces genes involved in resource allocation and transport, notably *SWEET12* (AT5G23660; ∼48-fold increase), a sucrose transporter required for efficient phloem loading (Chen et al., 2012), and *NAET1* (AT5G14940; ∼48-fold increase), a nicotianamine efflux transporter implicated in long-distance iron and copper transport (Chao et al., 2021). The induced gene set also reveals extensive regulatory reprogramming, with strong representation of non-coding RNAs, LINE-like retrotransposon transcripts (AT5G07215; ∼84-fold increase), and components of protein turnover and signaling, including RING/U-box E3 ubiquitin ligases (AT5G18260; ∼52-fold increase) and Ca^2^⁺-linked regulatory proteins (AT3G07490; ∼52-fold increase). Together, these changes point to coordinated activation of apoplastic remodeling, nutrient redistribution, and regulatory plasticity in response to ABA in *me1*.

Conversely, the strongest transcriptional repression in *me1* under ABA predominantly affects genes associated with defense-related functions, hormone and redox/ROS signaling, and cell-wall remodeling. Downregulated genes included *TSA1* (AT1G52410; ∼24-fold decrease), a JA-inducible Ca^2^⁺-binding factor required for wound-inducible ER-body formation and glucosinolate defense (Geem et al., 2019). *MYB28* (AT5G61420; ∼15-fold decrease), a transcriptional activator of aliphatic glucosinolate biosynthesis (Li et al., 2013), and a GH1 β-glucosidase (AT1G45191; ∼30-fold decrease) implicated in glucosinolate metabolism (Nakano et al., 2017). The redox/ROS-related components include H_2_O_2_ Response Gene 1 (AT2G41730; ∼12-fold decrease), which supports maintenance of root meristem activity under oxidative stress (Gong et al., 2021). This is accompanied by downregulation of two genes with established roles, FeSOD1 and a mitochondrial complex I subunit. Hormone-related regulators include *ARR11* (AT1G67710; ∼10-fold decrease), which is a type-B response regulator involved in cytokinin-controlled meristem/root growth (Hill et al., 2013) and *GASA9* (AT1G22690; ∼11-fold decrease), a gibberellin-regulated protein. The downregulated genes linked to altered cell-wall dynamics include a plasma-membrane monosaccharide transporter and *PME17* (AT2G45220; ∼8-fold decrease), a pectin methylesterase involved in homogalacturonan remodeling during root development (Sénéchal et al., 2014). Moreover, two strongly repressed RNA loci (each ∼60-fold downregulated), AT2G46572 and AT2G47015 (miR408, (Ma et al., 2015)), associated with laccase-related processes (Zhang et al., 2017a) further suggest that the

ABA response in this genotype involves a reprogramming of copper homeostasis and cell-wall phenolic oxidation pathways. As with the upregulated transcripts, the downregulated set was also enriched for non-coding RNAs, largely corresponding to pseudogenes and ribosomal RNA-related components.

In MS medium, the mutant shows enhanced expression of *ANAC060* (AT3G44290; 37-fold), which promotes seed germination by dampening sugar/ABA signaling by repressing *ABI5*, a central ABA-response regulator (Lopez-Molina et al., 2002). We also found reduced basal expression of genes at the intersection of stress/hormone signaling and seed-to-seedling programs. The most strongly downregulated transcripts include stress-associated regulators such as *ANAC019*, a transcriptional activator of JA signaling induced by ABA and abiotic stress (Tran et al., 2004), *ZAT10*, a repressor implicated in abiotic stress and JA signaling (Mittler et al., 2006), as well as seed-related factors including *MEE53* (Pagnussat et al., 2005), involved in embryo development, *RDO5*, involved in ABA signaling and regulator of seed dormancy (Xiang et al., 2014), and *LEA14*, a seed/stress dehydrin that contributes to longevity (Hundertmark et al., 2011). The set also contains a ribosomal structural protein and a pre tRNA (Leu), suggesting lowered investment in translation, alongside genes involved in glucosinolate breakdown and vesicle membrane fusion. Overall, the profile is consistent with unrestrained germination, but it likely predicts reduced robustness under environmental stress.

### Targeted KEGG enrichment of genotype-specific DEGs

To complement the MapMan analysis, we performed a targeted KEGG pathway enrichment focusing on the “Plant hormone signal transduction” pathway (Suppl. Fig. 7). Under control conditions, *me1* exhibits reduced expression of multiple components of ABA, ethylene, and jasmonate signaling, including *ABI1/ABI2* and other *PP2Cs* encoding genes, *EIN3*, *TCH4*, and the JA regulators *JAZ* and *MYC2*, whereas *DELLA* is the only consistently up-regulated factor, indicating a selective reconfiguration of hormone-responsive regulatory modules at baseline. Following ABA treatment, this pattern shifts markedly: *CPK28* and B-type *ARR* genes are down-regulated in *me1*, while A-type *ARRs*, *PIFs*, and *SAUR* genes are induced, reflecting stress-dependent rewiring of hormone signaling in the mutant.

### NADP-ME1 physically interacts with APX1 and MLP34

To examine whether NADP-ME1-mediated ABA responses in roots entail interactions with other proteins, we performed expression and co-immunoprecipitation (Co-IP) assays. First, we analyzed NADP-ME1 expression in roots of seedlings treated with 10 µM ABA. NADP-ME1 transcript levels were significantly upregulated, increasing ∼400-fold after 12h of ABA treatment, and pro*ME1*::GUS seedlings showed enhanced expression in roots (Suppl. Fig. 8A and B). We then used a construct for Co-IP assays in which an *ME1*::YFP fusion was expressed under the control of *NADP-ME1* promoter (p*ME1*::*ME1*::YFP). However, only a weak YFP signal was detected in the roots of ABA-treated seedlings (Suppl. Fig. 8C), and the protein levels were insufficient to allow reliable Co-IP assays.

As an alternative approach, we used seedlings expressing the pUBQ10::*ME1*::GFP construct (Fig. 1), in which GFP fluorescence was clearly detectable in roots (Suppl. Fig. 6D). Although *ME1*::GFP is constitutively expressed in these lines, we reasoned that ABA treatment would increase the availability of ABA-induced interaction partners, thereby enhancing the likelihood of detecting NADP-ME1-associated proteins. Consequently, Co-IP coupled with mass spectrometry identified several proteins as putative NADP-ME1 interactors (Fig. 8A; Suppl. Fig. 9; Suppl. Data 3).

**Figure 8.**
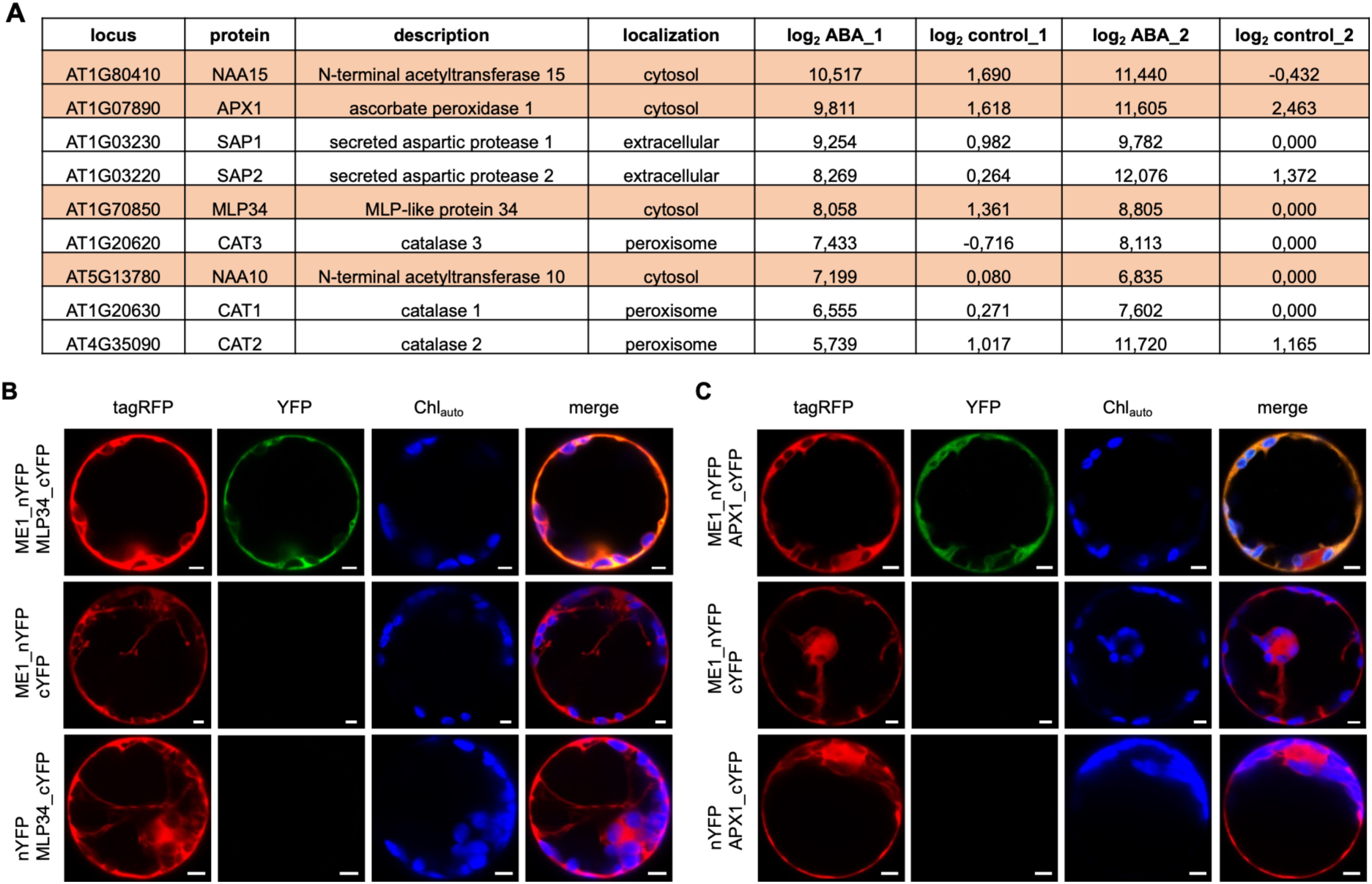
Validation of NADP-ME1 interactions with candidate proteins using BiFC assays. Bimolecular fluorescence complementation (BiFC) assays were performed by co-expressing AtNADP-ME1 fused to the N-terminal of YFP (nYFP) and candidate proteins fused to the C-terminal of YFP (cYFP) in a single 2-in-1 Gateway construct and expressing them in *N. tabacum* protoplasts. **A)** Specific proteins identified as putative interactors of NADP-ME1 via mass spectrometry. Proteins were selected based on enrichment under ABA treatment (Differences of log_2_ intensity > 5.6) and minimal detection under control condition (Differences of log_2_ intensity < 2.5) for the first replicate. All identified proteins are listed regardless of predicted subcellular localization. Proteins are presented in descending order of log_2_ intensity under ABA treatment for the first replicate (log_2_ABA_1). **B)** NADP-ME1 interacts with MLP34 in cytosol. Top row, Co-expression of ME1_nYFP and MLP34_cYFP showing YFP fluorescence. Second row, ME1_nYFP co-expressed with cYFP (negative control). Third row, nYFP co-expressed with MLP34_cYFP (negative control). **C)** NADP-ME1 interacts with APX1 in cytosol. Top row, Co-expression of ME1_nYFP and APX1_cYFP showing YFP fluorescence. Second row, ME1_nYFP co-expressed with cYFP (negative control). Third row, nYFP co-expressed with APX1_cYFP (negative control). tagRFP, red fluorescence; YFP, yellow fluorescence; Chl_auto_, chlorophyll autofluorescence; merge, overview of the protoplast. Scale bar: 5 µm.

We selected cytosol-localized candidates to validate their interaction with NADP-ME1 by Bimolecular Fluorescence Complementation (BiFC) in *Nicotiana benthamiana* protoplasts. The assay confirmed NADP-ME1 interactions with ascorbate peroxides 1 (APX1) and major latex-like protein 34 (MLP34), as YFP fluorescence was detected in cells co-expressing ME1-nYFP with APX1-cYFP or MLP34-cYFP, but not in the corresponding negative controls (Fig. 8B). Interactions with the other candidates were not observed by BiFC.

### Physiological contexts revealing a functional requirement for NADP-ME1

Tissue ABA concentrations can rise sharply as soil water content gradually declines (Zhang and Davies, 1989) or during salt stress (Jia et al., 2002). To position NADP-ME1 within a physiological context consistent with the ABA-and redox-associated functions identified in this study, we first monitored NADP-ME1 expression in *proME1*::GUS Arabidopsis seedlings subjected to progressive water deficit followed by re-watering. Under well-watered conditions, NADP-ME1 expression was very weak and confined to roots. As water deficit intensified, promoter activity increased progressively, reaching a maximum at the onset of visible wilting. At this stage, strong GUS staining was observed not only in roots but also in leaves. Upon re-watering, NADP-ME1 expression rapidly declined and returned to near-basal levels within two days (Fig. 9). These findings demonstrate that NADP-ME1 expression responds dynamically and reversibly to changes in plant water status. Despite this strong transcriptional induction of NADP-ME1 during water deficit, *me1* plants did not exhibit a visible drought-sensitive phenotype (Suppl. Fig. 10) or detectable alterations in photosynthetic performance (Suppl. Fig. 11). This suggests that NADP-ME1 is not essential for whole-plant survival under prolonged water limitation and that its loss may be buffered by redundant or compensatory pathways.

**Figure 9.**
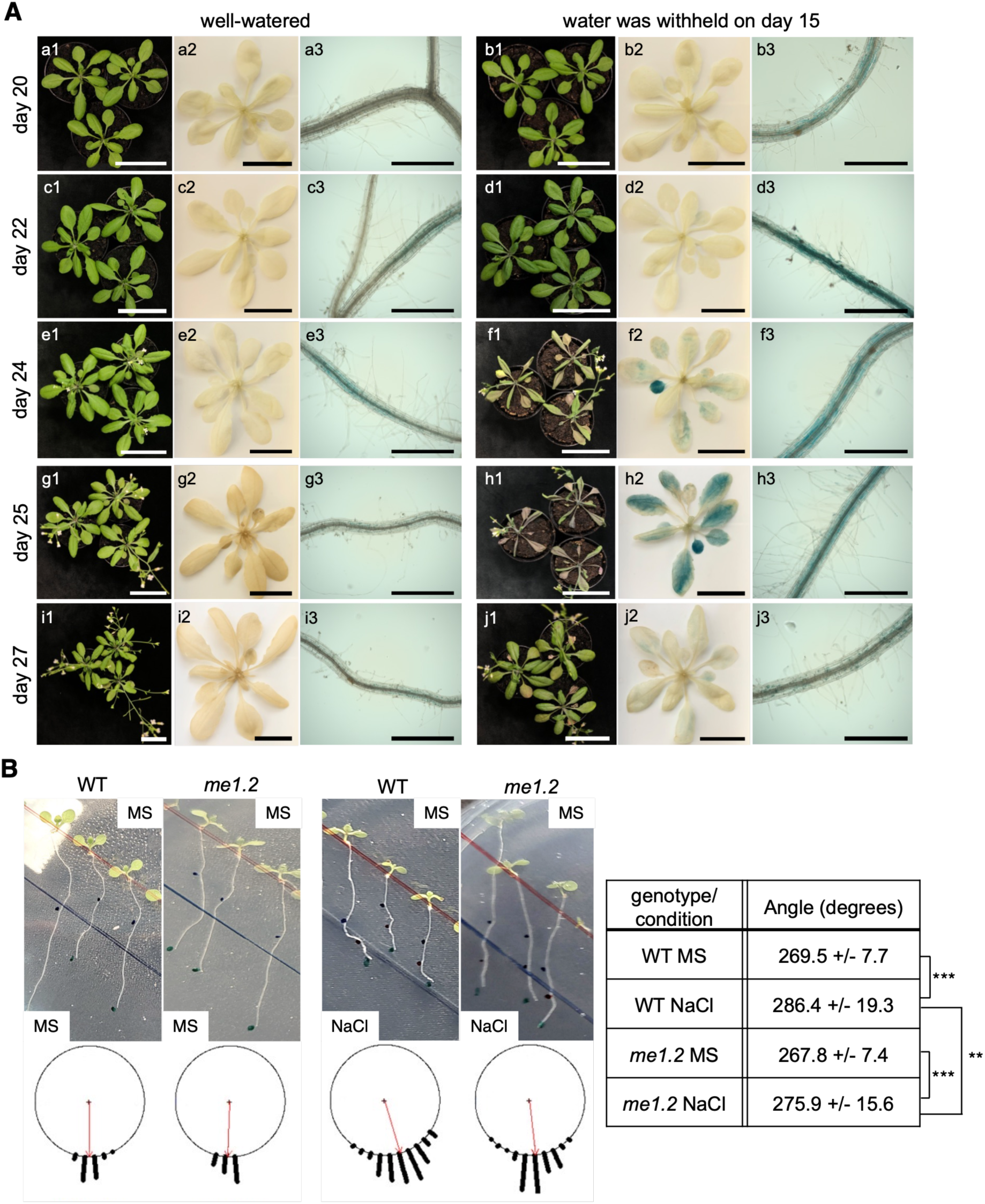
NADP-ME1 expression pattern during drought stress and halotropic root response under salt stress. **A)** Histochemical analysis of *NADP-ME1* promoter activity in well-watered and drought-stressed plants. Seedlings were grown for two weeks under well-watered conditions, after which watering was withheld. Wilting symptoms appeared after 10 days of drought treatment (day 24). On day 25 (one day post-wilting), plants were re-watered, and recovery was assessed two days later (day 27). a1-a3) 20-day-old seedlings under control conditions; b1-b3) 20-day-old seedlings under drought conditions; c1-c3) 22-day-old seedlings under control conditions; d1-d3) 22-day-old seedlings under drought conditions; e1-e3) 24-day-old seedlings under control conditions; f1-f3) 24-day-old seedlings under drought conditions; g1-g3) 25-day-old seedlings under control conditions; h1–h3) 25-day-old seedlings under drought conditions; i1-i3) 27-day-old seedlings under control conditions; j1-j3) 27-day-old seedlings after 2 days of re-watering. Scale bars: 5 cm in panels a1, b1, c1, d1, e1, f1, g1, h1, i1, j1; 2.5 cm in panels a2, b2, c2, d2, e2, f2, g2, h2, i2, j2; 500 µm in panels a3, b3, c3, d3, e3, f3, g3, h3, i3, j3. **B)** Halotropic response. Top: WT and *me1.2* seeds were germinated on vertically oriented plates containing ½ MS medium. After 6 days, part of the medium was replaced with ½ MS supplemented with 200 mM NaCl to induce a salt gradient. The red line marks the seed sowing line, and the blue line indicates the boundary between control medium and NaCl-supplemented medium. Dots represent the position of the root tip immediately before treatment, and 24 h and 48 h after treatment. Bottom: Radial histograms show the distribution root angles 48 h after treatment; the red arrow indicates the mean angle. In the table, data are presented as means ± SD (n = 42 for WT MS, 115 for WT NaCl, 39 for me1.2 MS, and 97 for me1.2 NaCl). Asterisks indicate significant differences in mean angles between genotypes, as determined by the Watson-Williams test (** *p* < 0.01; *** *p* < 0.001).

Because ABA partially mediates the evasive root response to elevated sodium chloride levels, enabling plants to minimize salt-induced damage (Geng et al., 2013), we next investigated the halotropic response of *me1.2* compared to wild type. Roots of the wild type exhibited a pronounced halotropic response, reorienting growth by approximately 16° away from the vertical axis. In contrast, *me1.2* roots showed a markedly reduced halotropic response, deviating by only ∼6° on average (Fig. 9B; Video 1), implicating NADP-ME1 in ABA-dependent salt avoidance.

## Discussion

The combined results indicate that NADP-ME1 acts as a redox regulator integrating ROS metabolism with ABA-controlled root growth. In wild-type plants, ABA suppresses primary root elongation and establishes an asymmetric auxin distribution at the root tip, consistent with previous reports that ABA-induced redox gradients drive localized auxin transport and growth arrest (Shkolnik-Inbar and Bar-Zvi, 2010; Sun et al., 2018; Pasternak et al., 2023). In contrast, *me1* mutants maintain partial root elongation under ABA, display bilateral auxin distribution, and accumulate superoxide, indicating that the enzyme contributes to proper redox balance required for ABA-mediated growth inhibition.

### NADP-ME1 coordinates ABA-induced ROS dynamics and hormone crosstalk in roots

NADP-ME catalyzes the oxidative decarboxylation of malate to pyruvate, generating NADPH that fuels antioxidant systems and helps maintain ROS homeostasis under stress conditions (Voll et al., 2012; Chen et al., 2019; Sun et al., 2019). Consistent with earlier reports, our data support a role for NADP-ME1 in shaping the root redox environment in the presence of ABA. Notably, the identification of APX1 as a NADP-ME1 interactor suggests a functional link between NADPH production and H_2_O_2_ detoxification, reminiscent of the APX1-centered antioxidant network described in Arabidopsis (Pnueli et al., 2003; Davletova et al., 2005). In line with this, *me1* roots accumulate O_2_^•⁻^ under ABA treatment, indicating a reduced redox buffering capacity. A similar imbalance between O_2_^•⁻^ and H_2_O_2_ has been observed in *apx1* mutants, where elevated ROS levels correlate with altered root development (Correa-Aragunde et al., 2013).

We also identified MLP34, a Bet v 1/PR-10-related protein, as an NADP-ME1-interacting partner. Members of the MLP family are thought to bind small hydrophobic ligands and have been implicated in stress-and hormone-associated signaling (Radauer et al., 2008; Fujita and Inui, 2021). Notably, MLPs are structurally related to the ABA receptor family (PYL/PYR), and MLP43 has been reported to promote ABA responses by interacting with SnRK2.6 and ABF1 and to enhance drought tolerance by limiting ROS accumulation, consistent with increased SOD and CAT activities (Wang et al., 2016). In this context, it is noteworthy that *SOD1* expression is more strongly downregulated in *me1* than in WT. Expression data from the Bio-Analytic Resource electronic Fluorescent Pictograph (BAR eFP) Browser, indicate that APX1 and MLP34 are constitutively expressed in roots (Winter et al., 2007), whereas NADP-ME1 is strongly induced by ABA. Under these conditions, interactions between NADP-ME1, APX1, and MLP34 may promote the formation of functional metabolons, facilitating substrate and product channeling or modulating the activities of associated enzymes.

Because redox homeostasis is a key determinant of root patterning and elongation (Tsukagoshi, 2016; Mhamdi and Van Breusegem, 2018), a shift in ROS composition provides a mechanistic explanation for the altered ABA response in *me1*. In wild-type roots, localized ROS accumulation promotes differential cell elongation, generating asymmetric auxin maxima and growth arrest on one side of the root tip. In *me1*, the bias toward superoxide likely disrupts this ROS gradient, leading to a more uniform auxin distribution and consequently reduced growth inhibition under ABA. The increased IAA content in *me1* roots under ABA treatment is consistent with redox-dependent regulation of auxin homeostasis (Pasternak et al., 2023). The ability of exogenous H_2_O_2_ to induce growth inhibition in both genotypes (Fig. 6B) further supports the view that in the context of ABA signaling, NADP-ME1 modulates root elongation primarily by controlling ROS levels.

ABA-dependent redox control is also well positioned to influence hormone crosstalk, because ROS are widely used as secondary messengers in multiple signaling pathways (Devireddy et al., 2021). Accordingly, ABA not only interacts with auxin but also with ethylene, gibberellin, and brassinosteroid signaling to fine-tune root growth (Qin et al., 2019; De Nittis et al., 2025). The observation that ethylene inhibitors restore the ABA-responsive phenotype further suggests that NADP-ME1-derived redox buffering normally constrains ethylene-dependent growth outputs under ABA. Together, these findings place NADP-ME1 at a convergence point where ABA-triggered ROS dynamics coordinate hormonal interactions to modulate root growth.

### NADP-ME1 modulates ABA-dependent transcriptional networks

Our transcriptomic analysis shows that NADP-ME1-dependent expression differences are minimal under control conditions but become strongly amplified upon ABA treatment. This supports a role for NADP-ME1 as a stress-conditional regulator that links cellular redox status with hormone signaling to fine-tune ABA-driven transcriptional programs and downstream root growth responses.

Consistent with elevated O_2_^•⁻^ detected in *me1* roots, the mutant shows reduced expression of *Fe-SOD*, encoding a O_2_^•⁻^-detoxifying enzyme contributing to ROS homeostasis in Arabidopsis roots (Alscher et al., 2002; Tsukagoshi, 2016) which supports root development (Dvořák et al., 2021). This suggests a diminished capacity to buffer O_2_^•⁻^, potentially linked to altered cytosolic NADPH availability. The redox disturbance is further reflected in the induction of canonical ROS/ABA-responsive modules, including defensins, cold-acclimation factors, and Hsp40/Hsp70 chaperones (Foyer and Noctor, 2005), indicating broader activation of stress-protective programs in the mutant background.

Beyond redox-associated changes, NADP-ME1 deficiency rewires hormone-linked transcriptional nodes that are central to growth control. Under ABA, *me1* displays enhanced induction of cytokinin-responsive A-ARR transcription factors together with repression of B-ARRs, shifting the ARR balance in a direction that could dampen ABA-mediated growth inhibition. This is notable given the well-established antagonism between cytokinin and ABA in regulating root growth (O’Brien and Benková, 2013; Guan et al., 2014), and the broader role of cytokinin-regulated transcription factors in modulating multiple hormonal pathways (Xie et al., 2018). In parallel, NADP-ME1 loss appears to weaken the GA-ABA growth-inhibition interface: PIF transcription factors (de Lucas and Prat, 2014) remain elevated during ABA exposure, while auxin-responsive growth regulators such as SAURs (Spartz et al., 2012) are not efficiently suppressed. This pattern is consistent with a model in which a PIF-SAUR module may contribute to sustaining cell elongation even under ABA by maintaining SAUR expression and counteracting growth arrest (Cañibano et al., 2025). Together, the sustained activity of PIF-and SAUR-dependent programs provides a mechanistic basis for the partial maintenance of root elongation in *me1* under ABA.

The transcriptome also points to an axis linking transcriptional rewiring to the altered auxin-redistribution phenotype. The upregulation of *RG1* and *LZY1*, implicated in gravitropic auxin signaling (Taniguchi et al., 2017; Jiao et al., 2021), offers a plausible connection between hormone signaling, directional auxin transport, and the *DR5* distribution changes observed in *me1* roots under ABA. Thus, NADP-ME1 loss not only perturbs ROS buffering but also shifts hormone-regulated transport and growth-control circuits that determine spatial auxin patterning at the root apex.

Importantly, in our analysis several of the strongest genotype-dependent ABA-regulated genes fall outside statistically enriched MapMan bins. Their known functions nevertheless align closely with the physiological and cellular phenotypes, underscoring the complementarity of bin-based enrichment and targeted gene-level interpretation, and highlighting the value of highly regulated “unassigned” genes in complex stress responses.

Taken together, ABA triggers a broadly conserved core response in both genotypes, whereas NADP-ME1 selectively fine-tunes a subset of redox-, hormone-, and growth-associated pathways. In the absence of NADP-ME1, the balance of these outputs shifts toward growth-promoting signaling and altered auxin transport, biasing the response toward root elongation rather than full ABA-induced growth arrest. Importantly, however, *me1* roots under ABA still do not attain the elongation rates seen in WT and *me1* under control conditions, indicating that ABA activates additional growth-inhibitory mechanisms that constrain elongation independently of NADP-ME1.

### Integrative model

Overall, our data support a model in which cytosolic NADP-ME1 couples ABA signaling to local redox control at the root apex, thereby tuning ABA-driven transcriptional reprogramming and limiting root elongation under stress. In roots of WT plants grown on MS medium, *NADP-ME1* expression is barely undetectable, whereas ABA strongly induces its expression (Suppl. Fig. 8A, B; (Arias et al., 2018). We therefore propose that, following ABA induction, NADP-ME1 acts as part of a root-tip redox module together with APX1 and MLP34 to regulate the balance between O_2_^•⁻^ and H_2_O_2_ (Fig. 10). In wild type, increased NADP-ME1 activity at the root tip boosts the cytosolic NADPH pool, thereby promoting ascorbate recycling and sustaining APX1-dependent H_2_O_2_ turnover. In parallel, superoxide dismutase-mediated conversion of O_2_^•⁻^ to H_2_O_2_ generating a spatially confined redox signal at the root tip. We propose that this redox signal is required to establish asymmetric auxin distribution and, consequently, to inhibit root growth. Consistently, H_2_O_2_ has been shown to impair PIN2 vesicle trafficking by disrupting cytoskeletal function (Zwiewka et al., 2019). Moreover, Arabidopsis *apx1* mutants have increased H_2_O_2_ accumulation in roots and shorter roots (Correa-Aragunde et al., 2013). In *me1*, reduced NADPH availability likely limits ascorbate regeneration, compromising APX1 activity and shifting ROS speciation toward O_2_^•⁻^ accumulation. This change would flatten the root-tip redox gradient, disrupt ABA-dependent auxin asymmetry, and thereby permit partial root elongation despite ABA exposure. Together, this model positions NADP-ME1 as an integrator of ABA signaling, ROS processing, and auxin patterning, linking cytosolic redox metabolism to hormone crosstalk that determines root growth outcomes under stress.

**Figure 10.**
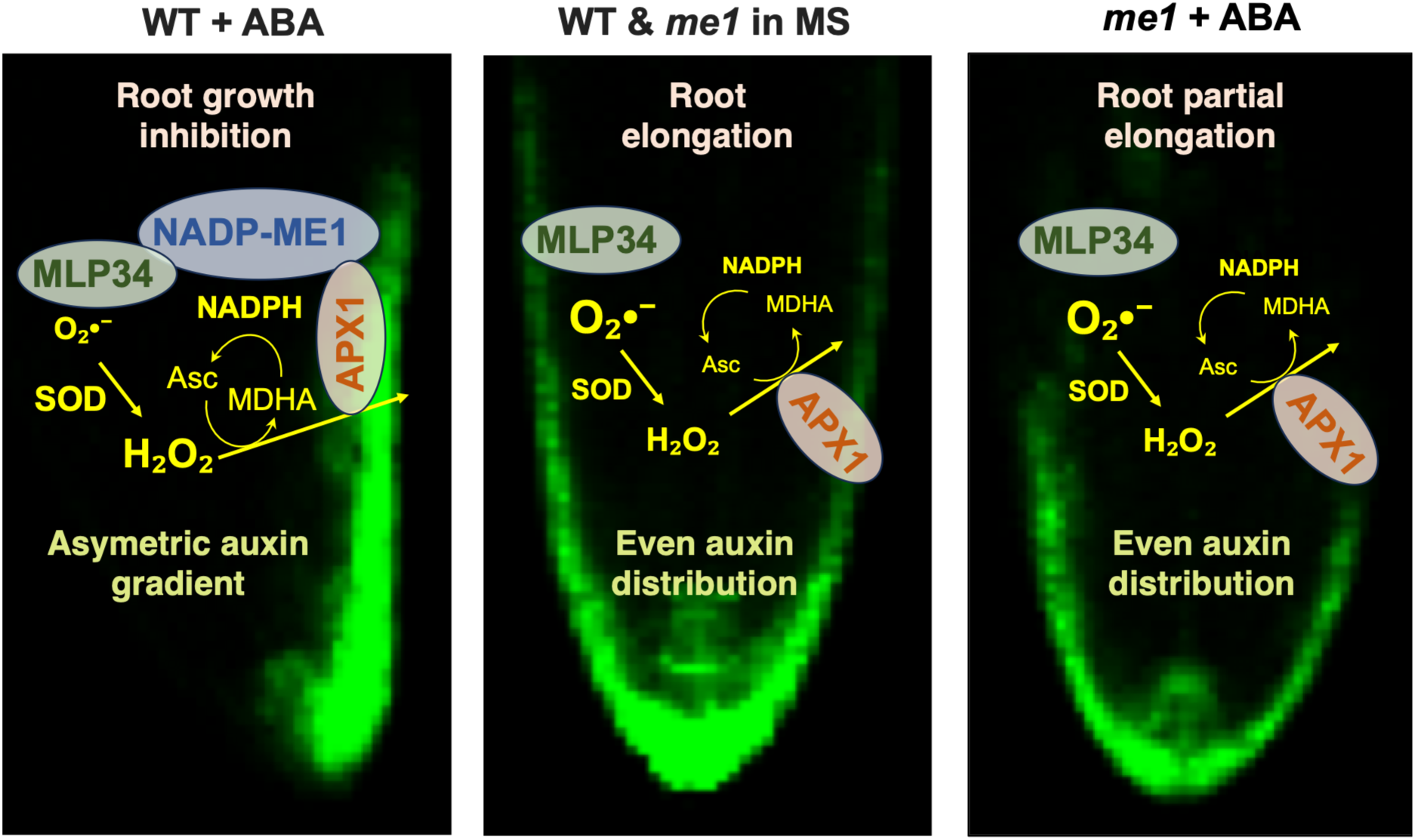
Proposed model for NADP-ME1 function at the root apex. Root elongation of WT and *me1* seedlings is similar on MS medium, consistent with the barely undetectable expression of NADP-ME1 in WT under these conditions (Suppl. Fig. 8A, B; (Arias et al., 2018)). In the presence of ABA, however, NADP-ME1 is induced in WT roots and is proposed to function within a local redox module involving APX1 and MLP34. In this context, NADP-ME1 supports ROS homeostasis and helps maintain the balance between O_2_^•⁻^ and H_2_O_2_, thereby contributing to asymmetric auxin gradients and growth inhibition in wild-type roots. In *me1*, the absence of NADP-ME1 is proposed to impair ROS homeostasis, resulting in altered ROS gradients and a more uniform auxin distribution, consistent with reduced ABA-induced inhibition of root growth.

### ABA-linked NADP-ME1 function in adaptation to drought and salt stress

ABA acts as a central integrator of drought and salinity signaling, reshaping root system architecture by maintaining axial growth while restricting the initiation and expansion of lateral organs, thereby enhancing soil foraging efficiency under water-limiting conditions (Sharp et al., 2004; Deak and Malamy, 2005; De Smet et al., 2006; Dinneny, 2019). ABA accumulation is required to sustain primary root elongation at low water potentials (Saab et al., 1990), whereas lateral root formation is progressively repressed as water availability declines, a process in which ABA plays a key regulatory role (Deak and Malamy, 2005; De Smet et al., 2006). In parallel, salt stress perturbs auxin homeostasis and distribution in primary roots (Warnecke et al., 2009; Tessi et al., 2023), and ROS-auxin crosstalk further modulates this response through PIN-mediated auxin transport (Fu et al., 2019). Together, hormonal and redox signals converge to regulate adaptive root growth.

Within this framework, our results identify NADP-ME1 as a component of the ABA-responsive redox machinery that supports root plasticity under water deficit and salinity. NADP-ME1 expression is strongly induced during progressive drought and levels are declines upon rewatering, consistent with established ABA dynamics under drought stress. We hypothesize that NADP-ME1 contributes to cytosolic redox homeostasis by supplying NADPH, a function that may be particularly important in spatially restricted cellular contexts. The absence of a pronounced drought phenotype suggests functional redundancy or metabolic compensation, possibly through other NADP-dependent dehydrogenases or alternative NADPH-generating systems. Future work aimed at dissecting this redundancy will be essential to define the quantitative contribution of NADP-ME1 to cytosolic NADPH pools under stress. Consistent with this role, *me1* mutant roots exhibit a markedly attenuated halotropic response. Because halotropism requires coordinated ABA signaling, ROS dynamics, and auxin redistribution at the root apex (Wang et al., 2009; Galvan-Ampudia et al., 2013; Geng et al., 2013), the reduced root reorientation observed in *me1* supports a model in which NADP-ME1-derived NADPH production maintains localized redox homeostasis required for proper auxin patterning and directional growth. These findings place NADP-ME1 at the intersection of ABA signaling, cytosolic redox control, and auxin transport, fine-tuning adaptive root growth under stress.

Drought and salinity are major environmental constraints that reduce water availability and impose osmotic stress on plants. The convergence of these conditions on ABA signaling and redox remodeling highlights the importance of metabolic systems, such as NADP-ME1-dependent NADPH production, in maintaining cellular homeostasis while enabling adaptive root growth. Efficient acquisition of water and nutrients is critical for plant productivity, particularly under increasingly variable environmental conditions. Thus, a deeper understanding of the metabolic and signaling networks controlling root development, including the ABA-redox-auxin interface described here, will facilitate the rational design of strategies to improve stress resilience and sustain crop yields in the face of climate change.

## Material and method

### Plant lines and growth conditions

In this study, we used *Arabidopsis thaliana* ecotype Columbia-0 (Col-0) as wild-type (WT) and T-DNA insertion lines of *NADP-ME1* (AT2G19900), namely *me1.1* (SALKseq_047029.1), *me1.2* (SALKseq_036898.0), and *me1.3* (SALKseq_137251.1). *A. thaliana* T-DNA insertion lines were obtained from the Nottingham Arabidopsis Stock Centre (NASC, http://arabidopsis.info/). All T-DNA insertion lines were screened via PCR with specific genomic and T-DNA left border primers (Suppl. Table 1), and homozygous knockout mutants were isolated for all lines. Reporter lines pro*ME1*::GUS was screened previously (Gerrard Wheeler et al., 2005). The TCSn:GFP and DR5:RFP reporter lines were produced in our previous work (Tessi et al., 2023). The double reporter-mutant combinations (TCSn:GFP/*me1.2*, TCSn:GFP/*me1.3*, DR5:RFP/*me1.2*, and DR5:RFP/*me1.3*) and reporter-WT lines (TCSn:GFP/WT and DR5:RFP/WT) were generated through genetic crosses. The p*ME1*::*ME1*::YFP and p*ME1*::YFP reporter lines were produced in our previous work (Arias et al., 2018). For plants growing in soil, seedlings were grown in *Floradur® B Seed* substrate under a 16 h light / 8 h dark photoperiod at 22 °C (day) / 18 °C (night) in a controlled growth chamber with a photosynthetic photon flux density (PPFD) of 120 µmol m⁻^2^ s⁻^1^. For plants growing in plates, seeds were sterilized in 70% (v/v) ethanol for 5 min, followed by 100% ethanol for 1 min, then rinsed three times with sterile double-distilled water. Sterilized seeds were sown on half-strength Murashige and Skoog (½ MS; M0221, Duchefa Biochemie) medium (Murashige and Skoog, 1962) solidified with 1% (w/v) agar (AGA03, FORMEDIUM). Seeds were stratified at 4 °C in darkness for 72 h before transfer to the growth chamber under the same environmental conditions described above.

### Root length measurement

Root length measurement of WT and mutant lines (*me1.2* and *me1.3*) were performed on sterile ½ MS medium with or without 1 µM abscisic acid (ABA; 90769, Sigma-Aldrich). ABA was dissolved in dimethyl sulfoxide (DMSO; A994.1, Carl Roth). Twelve seeds per line were sown per plate in randomized order, with six technical and three biological replicates per treatment. Images were taken using a Canon EOS 2000D camera, and root length was measured using FIJI (Schindelin et al., 2012). For the hormone inhibitor-based analyses, five inhibitors were incorporated into the growth medium, with or without 1µM ABA. These included the ethylene biosynthesis inhibitor L-α-(2-Aminoethoxyvinyl)-glycine hydrochloride (AVG-HCl; 1 µM; A6685, Sigma-Aldrich), the ethylene perception inhibitor silver nitrate (AgNO₃; 10 µM; No. 2246.2, Carl Roth), the auxin efflux inhibitor 2,3,5-triiodobenzoic acid (TIBA; 5 µM; T5910, Sigma-Aldrich), the auxin influx inhibitor 3-chloro-4-hydroxyphenylacetic acid (CHPAA; 20 µM; 224529, Sigma-Aldrich), and the Ca^2^⁺ channel inhibitor lanthanum chloride (LaCl_3_; 100 µM; 298182, Sigma-Aldrich). Five biological replicates were included for inhibitor experiments. For hydrogen peroxide (H_2_O_2_) treatment, seedlings were sown on the ½ MS medium with or without 1 µM ABA in combination with H_2_O_2_ (1 mM or 2 mM). Four biological replicates were included. Growth conditions and root length measurements were as described above.

### RNA isolation, reverse transcription, and qPCR analysis

Total RNA of root from WT and mutants (*me1.1*, *me1.2*, and *me1.3*) was isolated using the EURX Universal RNA Purification Kit (E3598, Roboklon GmbH, Germany). For RNA-seq, total RNA was isolated using a modified SDS-LiCl method (Vennapusa et al., 2020). The quality and concentration were tested and measured with the NanoDrop One UV-Vis spectrophotometer (Thermo Fisher scientific) and gel electrophoresis. After the removal of possible genomic DNA contaminations with an Ambion DNA-free DNA removal kit (AM1906, Thermo Fisher scientific), 750 ng RNA was transcribed into the complementary DNA (cDNA) with RevertAid reverse transcriptase kit (K1691, Thermo Fisher scientific) using an oligo(dT) primer (Suppl. Table 1) according to the manufacturer’s instructions. cDNA levels were normalized to *Actin2* (AT3G18780) as an endogenous control (Czechowski et al., 2005). Transcripts abundance for mutants was measured via RT-PCR using specific primers (Suppl. Table 1).

Transcript analysis of mutants via qPCR was performed with KAPA SYBR FAST qPCR Master Mix (KK4618, Sigma-Aldrich) and 2 µL of 1:10 diluted cDNA in a CFX96 C1000 Touch Real-Time PCR instrument (Bio-Rad, USA) according to the manufacturer’s instructions. Specific primers for qPCR were designed using Primer-Blast (NCBI) and snapgene (Suppl. Table 1). The expression of differential *NADP-ME1* was analyzed and expressed using the ΔΔCT method (Schmittgen and Livak, 2008).

For transcript analysis of *NADP-ME1* under ABA treatment, WT was grown on ½ MS medium for 10 days and then transferred to fresh ½ MS medium with or without 10 µM ABA for 12 h. Root samples were collected for qPCR using the same method as described above, with four biological replicates per treatment.

### Plasmid construction and plant transformation

The coding sequence of the *NADP-ME1* were amplified from the cDNA of WT root using HF buffer and Phusion polymerase (F530L, Thermo Fisher scientific) and specific primers (Suppl. Table 1). Then, the amplified fragment was sub-cloned into pCR-TOPO-Blunt (TOPO, 450245, Thermo Fisher scientific). The TOPO vector was used as template for further PCR-dependent fragment constructions according to Gibson cloning method (Gibson et al., 2009). Primers for Gibson cloning were designed with a 20-bp overlap with the binary vector (Suppl. Table 1).

For the subcellular localization, the vector pUBQ10::GFP was used to construct the C-and N-terminal GFP-fusions (Kunz et al., 2014). Vectors linearized with BamHI/SacI were used in a Gibson assembly reaction with the amplified *NADP-ME1* coding sequence to yield in-frame GFP fusions, whose expression was under the control of the constitutive ubiquitin-10 (UBQ10, AT4G05320) promoter. The vector pUBQ10::*ME1*::GFP constructed for the expression of C-terminal fusion proteins as well as the vector pUBQ10::GFP::*ME1* constructed for the expression of N-terminal fusion proteins were transformed into *Agrobacterium tumefaciens* (strain GV3101 with pMP90) and selected on Yest Extract beef medium (YEB) plates containing rifampicin (100 mg/mL), gentamycin (25 mg/mL), and kanamycin (50 mg/mL). *Agrobacterium tumefaciens* with the target construct was transformed into Col-0 by flower dip (Clough and Bent, 1998).

For biomolecular fluorescence complementation (BiFC), the vectors constructed using the 2in1 cloning system were used for BiFC analyses (Grefen and Blatt, 2012). The insertion of the *NADP-ME1* amplicons into pDONR221 P2R-P3 entry vector carrying the attP2 and attP3 recombination sites was accomplished by BP reactions using the BP Clonase TM II enzyme mix (11789020, Thermo Fisher Scientific). The same enzyme was used for insertion of interaction candidates into pDONR221 P1-P4 carrying the attP1 and attP4 recombination sites. Subsequently, entry clones were integrated with the corresponding destination vector pBiFCt-2in1-CC. The reaction was catalyzed by LR Clonase TM II Mix (11791020, Thermo Fisher Scientific). Finally, the expression constructs containing the *NADP-ME1* coding sequence without stop codon (-TGA) C-terminally fused to the N-terminal part of YFP (NADP-ME1 (-TGA)_nYFP) and the coding sequence of the interaction candidates coding sequence without stop codon (-TGA) C-terminally fused to the C-terminal part of YFP (interaction candidate (-TAG)_cYFP) were generated.

### Auxin and cytokinin distribution detection

Auxin and cytokinin distribution were examined in seedlings of reporter-WT (DR5:RFP/WT and TCSn:GFP/WT) and reporter-mutant (DR5:RFP/*me1.2*, DR5:RFP/*me1.3,* TCSn:GFP/*me1.2*, and TCSn:GFP/*me1.3*) grown on ½ MS medium with or without 1 µM ABA for 4 days. Auxin and cytokinin distribution were visualized using an TCS SP8 confocal laser scanning microscope (CLSM, Leica, Germany). Each assay was independently repeated at least three times. For auxin signal detection, RFP fluorescence was excited at 552 nm, and emission was collected between 579 and 633 nm. For cytokinin signal detection using the TCSn reporter system, GFP fluorescence was excited at 488 nm and emission was detected at 499 to 520 nm. Chlorophyll autofluorescence was detected at 686-711 nm. Propidium iodide (PI; 537059, Sigma-Aldrich) staining was employed to visualize cell wall in roots (Zhang et al., 2017b). Briefly, roots were incubated in PI staining solution for 5 min, rinsed with distilled water, and then visualized under CLSM. PI fluorescence was excited at 488 nm, and emission was collected between 600 and 650 nm.

### Subcellular localization

Infiltration of Agrobacterium cultures of pUBQ10::GFP, pUBQ10::*ME1*::GFP, and pUBQ10::GFP::*ME1* constructs into 4-week-old wild tobacco leaves (*Nicotiana benthamiana*) separately and protoplast isolation were performed according to Waadt and Kudla (2008). In briefly, Positive Agrobacterium clones were cultured in YEB medium with respective antibiotics until OD600 reach 1. Then, cells were resuspended in 10 mM MgCl_2_, 10 mM 2-morpholinoethanesulfonic acid (MES), and 100 μM acetosyringone (AS), and infiltrated into tobacco leaves using a syringe. After 3d infiltration, infiltrated leaf protoplasts were isolated by isolation buffer (20 mM MES-KOH pH 5.6, 10 mM CaCl_2_, 20 mM KCl, 0.1 % BSA (w/v), 0.3 % macerozym (w/v), 1 % cellulase (w/v), and 0.4 M mannitol), Pellet protoplasts by centrifugation 100 g for 3 min, discard the supernatant. Then the protoplast pellet was resuspended in W5 buffer (154 mM NaCl, 125 mM CaCl_2_, 5 mM KCl, 2 mM MES-KOH pH 5.6) for further imaging. Fluorescence was monitored under CLSM. GFP fluorescence and chlorophyll autofluorescence were detected as described above.

### PEG-mediated transformation in protoplast for BiFC

PEG-mediated transformation was performed following a modified protocol from Yoo et al. (2007). *Nicotiana benthamiana* were grown in soil for at least 3 weeks. Young leaves were cut into strips and digested overnight in enzyme solution (0.5% Cellulase R10, 0.15% Macerozyme R10, 10 mM CaCl₂, 0.4 M mannitol, and 5 mM MES, pH 5.6) at room temperature in the dark. The resulting mixture was filtered through a 90 µm mesh, and protoplasts were collected by low-speed centrifugation and purified using flotation buffer (10 mM CaCl_2_·2H_2_O, 0.2 mM KH_2_PO_4_, 1 mM KNO₃, 1 mM MgSO_4_·7H₂O, 5 mM MES, 1 mM PVP-10, and 0.46 M sucrose, pH 5.6) and W5 buffer washes. Purified protoplasts were resuspended in MaMg solution (0.4 M mannitol, 15 mM MgCl_2_, and 20 mM MES, pH 6.5) and mixed with 50 µg plasmid DNA (ME1_nYFP and candidate_cYFP 2in1 construct). Transformation was induced with 40% (w/v) PEG 1500 for 20-30 min in darkness. After PEG transformation, protoplasts were washed with W5 buffer, centrifuged, resuspended, and incubated overnight at 22 °C in the dark before imaging. Fluorescence signals were visualized by CLSM, tagRFP fluorescence and chlorophyll autofluorescence were detected as described above. YFP fluorescence was excited at 488 nm, and emission was collected between 517 and 553 nm.

### Phytohormone quantification by LC-MS

Seedling of the WT as well as mutant lines (*me1.2* and *me1.3*) were grown on ½ MS medium with or without 1 µM ABA and harvested after 48 h. Seven biological replicates with approximately 50 mg of plant material each were used. Phytohormones were extracted in a two-phase solvent-solvent extraction using isopropanol/water/HCl and dichloromethane, following a protocol adapted from Pan et al. (2010). Phytohormones extraction from germinating seeds was performed as described in Wewer et al., 2026 (unpublished), with an additional clean-up step to remove excessive triacylglycerol (TAG) from the samples prior to injection. To this end, dried phytohormone extracts were dissolved in 500 µl methanol and 500 µl of cold (4 °C) hexane were added. Samples were vortexed and centrifuged for 5 minutes at 10.000 g for phase separation. The upper hexane layer containing TAG was carefully removed and discarded. Another 500 µl of hexane were added and the procedure was repeated. The remaining 500 µl methanol phase were dried once again under a stream of nitrogen and phytohormones were re-dissolved in 50 µl methanol and 50 µl water for LC-MS analysis. Phytohormone separation was done on a C18 XSelect™ HSST3 HPLC column (2.5 µm, 3.0 mm x 150 mm, 100 Å, (Waters)) at a flow rate of 0.35 mL/min using a binary gradient. Solvent A was H_2_O + 0.1 % (v/v) formic acid, solvent B was methanol + 0.1% (v/v) formic acid. The gradient started at 2% B, increasing from 2% B to 99% B from 1 min to 18 min, held at 99% B for 10 minutes and decreased to 2% B at 30 min. This was held for another 5 minutes for equilibration.

Phytohormone analysis by LC-MS was performed on a QTrap 5500 QQQ instrument (Sciex) using multiple reaction monitoring (MRM) as described in Wewer et al., 2026. The analysis was done in fast polarity switching mode (positive and negative mode).

Phytohormones were quantified using calibration curves based on area ratios in relation to stable isotope-labelled internal standards (I.S.) as described in Wewer et al., 2026. Specifically, IAA and conjugated IAA were quantified in relation to the I.S. ^2^H_5_-IAA. JA, JA-Ileu and OPDA were quantified in relation to the I.S. ^2^H_6_-JA; tZ, cZ, IP and IPR were quantified in relation to the I.S. ^2^H_5_-tZ, ^15^N_4_-cZ, ^15^N_4_-IP and ^2^H_6_-IPR., tZ7G/tZOG and cZOG/tZ9G were quantified in relation to the I.S. ^2^H_5_-tZ7G/^2^H_5_-tZOG and ^2^H_5_-tZ9G. No stable isotope-labelled internal standards were available for tZR and cZR, which were quantified based on their peak area relative to external calibration curves.

### ROS Detection via NBT Staining Assays

Seedlings of WT and mutant lines (*me1.2* and *me1.3*) were grown on ½ MS medium with or without 1 µM ABA for 4 days and subsequently collected for ROS detection. Nitro blue Tetrazolium (NBT, N6639, Sigma-Aldrich) staining was used to detect superoxide (O_2_•⁻) in situ, as previously described by Dunand et al (2007). Seedlings were incubated with 2 mM NBT in 20 mM K-phosphate with 0.1 M NaCl at pH 6.1 for 1 h, and then transferred to distilled water. Root tips were imaged under bright-field illumination using Zeiss Axiolab 5 microscopy. Approximately 20 roots per genotype were imaged, with three biological replicates.

### Co-immunoprecipitation and Western Blot analysis

For co-immunoprecipitation (Co-IP) analyses, pUBQ10::*ME1*::GFP and pUBQ10::GFP seedlings were grown on ½ MS medium for 10 days and then transferred to fresh ½ MS medium with or without 10 µM ABA for 12 h. Fluorescence was observed using CLSM. Roots were stained with PI as described above, GFP and PI fluorescence were detected following the same procedures. Root tissues were subsequently collected for Co-IP assays. Two biological replicates were included per treatment. Approximately 150 mg of root tissue was ground in liquid nitrogen and homogenized in 1 ml lysis buffer (50 mM Tris-HCl, pH 7.5; 150 mM NaCl; 10 mM MgCl_2_; 0.5 mM EDTA; 1 mM DTT; 1% Triton X-100; 1 mM PMSF). After 1 h incubation on ice with occasional vortexing, cell debris was removed by centrifugation (15 min, 4 °C, maximum speed). The supernatant was transferred to a new tube containing 25 µl equilibrated anti-GFP agarose beads (AB_2631357, GFP-Trap® Agarose, ChromoTek) and incubated overnight at 4°C with rotation. Beads were washed several times with lysis buffer, and bound proteins were eluted twice with 80 µl acidic elution buffer (200 mM glycine, pH 2.5) and neutralized with 1 M Tris (pH 10.4). 10 µl of the elution was used for western blot analysis, and the remaining eluates were stored at-80 °C for mass spectrometry.

For western blotting, protein fractions were mixed with 2x Laemmli buffer (120 mM Tris-HCl pH 6.8, 20% glycerol, 4% SDS, 0.04% (w/v) bromophenol blue, 10% β-mercaptoethanol), heated for 5 min at 95°C, and separated on a 10% sodium dodecyl sulfate–polyacrylamide gel electrophoresis (SDS-PAGE). Proteins were transferred to a nitrocellulose membrane and detected using anti-GFP antibody (11814460001, Sigma-Aldrich; 1:1000) and AP-conjugated secondary antibody (31320, Thermo Fisher Scientific; 1:5000). Signal was developed using 5-bromo-4-chloro-3-indolyl-phosphate (BCIP, No. 6368.1, Carl Roth)/NBT substrate, and images were captured using a cell phone.

### Proteomic mass spectrometry analysis

Eluates from GFP-Trap® Agarose were prepared for mass spectrometric analysis by in-gel digestion with trypsin essentially as described earlier (Grube et al., 2018). Briefly, eluates were shortly separated in a polyacrylamide-gel, proteins reduced with dithiothreitol, alkylated with iodoacetamide and digested overnight with 0.1 µg trypsin. Peptides were extracted from the gel, dried in a vacuum concentrator, resuspended in 17 µl 0.1 % (v/v) trifluoroacetic acid and analyzed by liquid chromatography coupled mass spectrometry as described with modifications mentions below (Prescher et al., 2021). First, peptides were trapped on a 2 cm long precolumn and then separated by a one-hour gradient on a 25 cm long C18 analytical column using an Ultimate 3000 rapid separation liquid chromatography system (Thermo Fisher Scientific). Second, peptides were injected in a Fusion Lumos (Thermo Fisher Scientific) mass spectrometer online coupled by a nano-source electrospray interface. The mass spectrometer was operated in data-dependent positive mode. Peptide and protein identification were carried out with MaxQuant version 2.5.2.0 (Max Planck Institute for Biochemistry, Germany) with standard parameters if not stated otherwise. Protein sequences from *Arabidopsis thaliana* (downloaded from “The Arabidopsis Information Resource” (TAIR): Araport11_pep from 20220914) were used for searches with standard parameters. Identified proteins were filtered: contaminants, “identified by site” proteins, reverse hits and proteins only identified with one peptide and with no quantitative values were removed.

### Expression analysis of *AtNADP-MEs* under drought stress and ABA treatment

Two-week-old seedlings of pro*ME1*::GUS were used to perform the drought stress by water withdrawal. After 10-day stop watering (day 24), the seedlings started to wilt. On day 25, the plants were watered again. The seedlings recovered 2d after re-watering (day 27). Whole seedlings were harvested on days 20, 22, 24, 25, and 27 and subsequently immersed in GUS staining solution (100 mM phosphate buffer (pH 7.0), 2 mM potassium-ferrocyanide, 2 mM potassium-ferricyanide, 0.1% Triton X-100, 10 mM ethylenediaminetetraacetic acid (EDTA) and 1 mM X-Gluc) (Jefferson, 1987) using the vacuum infiltration method and then incubated at 37°C for overnight. After incubation, seedling was transferred to 70% ethanol solution to bleach the chlorophyll and it was then observed under the stereo microscope (Nikon SMZ800) and cell phone. Three biological replicates were included. Five-and Ten-day-old pro*ME1*::GUS seedlings were grown on ½ MS medium for 10 days, then transferred to ½ MS medium with or without 10 µM ABA for an additional 12 hours. Seedlings were then collected and subjected to GUS staining solution as described above. Roots were observed using Zeiss Axiolab 5 microscope as described as above. Ten-day-old pUBQ10::GFP, pUBQ10::*ME1*::GFP, p*ME1*::YFP, and p*ME1*::*ME1*::YFP seedlings were similarity treated with 10 µM ABA for 12 hours. Roots were stained with PI as described above, and GFP, YFP, and PI fluorescence were detected following the same procedures. Phenotyping of *me1* mutants (*me1.2* and *me1.3*) under drought stress. Two-week-old seedlings of WT and *me1* mutants were subjected to drought stress by withholding water. After two weeks of water deprivation (day 28), the seedlings began to show visible wilting symptoms. On day 29, plants were re-watered to assess their recovery response. Photographs were taken by a cell phone, and photosynthetic performance was measured using Pulse-Amplitude-Modulation fluorometry (Schreiber, 2004).

### Halotropic response assay

WT and *me1.2* seeds were sown on ½ MS medium and plates were kept in a vertical position during growth. After 6 days, a portion of the medium was replaced with ½ MS supplemented with 200 mM sodium chloride to create a salt gradient. The cutting edge of the replaced medium was positioned 1 cm from the sowing line. Root growth was monitored, and the angle of each root was measured 48 h after treatment using Fiji. Root bending angles were analyzed as circular data.

### Transcriptomic analysis

Transcriptomic analysis was performed on the WT and *me1.2* seedlings grown on ½ MS medium with or without 1 µM ABA. To ensure developmental stage-matched comparisons despite ABA-induced growth differences, *me1.2* seedlings were harvested after 48 h of ABA treatment, whereas wild-type seedlings were collected after 80h; under control conditions, both genotypes were harvested at 80h. Four biological replicates were analyzed per treatment. Total RNA was extracted as described above, and libraries were prepared using the Lexogen QuantSeq 3′ mRNA-Seq Library Prep Kit and sequenced on an Illumina NovaSeq 6000 platform, generating ∼10 million 1×100 bp single-end reads per sample.

Raw read quality was assessed with FastQC, and adapter trimming and quality filtering were conducted using fastp (v0.23.2), followed by quality reassessment with MultiQC. Reads were aligned to the *Arabidopsis thaliana* TAIR10 genome using HISAT2 (v2.2.1) with splice-aware mapping (--dta), and alignments were processed and validated with samtools (v1.12). Gene-level counts were obtained using featureCounts (v2.0.3) with forward-stranded read assignment. Mapping quality, read distribution, and replicate consistency were verified through coverage visualization in IGV and correlation heatmaps generated from DESeq2-normalized counts (Suppl. Fig. 12). Differential expression analysis was carried out in DESeq2 (v1.40.2) using a two-factor design (genotype × treatment), testing four contrasts: (1) WT ABA vs WT control, (2) *me1.2* ABA vs *me1.2* control, (3) *me1.2* ABA vs WT ABA, and (4) *me1.2* control vs WT control. Genes with adjusted *p* < 0.05 and |log_2_FC| > 1 were considered differentially expressed. PCA and heatmap analyses confirmed replicate clustering and treatment effects.

### Targeted KEGG pathway enrichment

Targeted pathway enrichment analysis was performed using the KEGG pathway *Plant hormone signal transduction – Arabidopsis thaliana* (ath04075) via the KEGG Mapper platform. Significantly differentially expressed genes (DEGs) identified from pairwise comparisons between *me1* mutants and wild type under control and ABA treatment were used as input. Up-regulated and down-regulated gene sets were analyzed separately to assess direction-specific changes in hormone signaling components. Enrichment was evaluated based on the presence and distribution of DEGs within the curated KEGG pathway map.

### MapMan enrichment

Functional enrichment of MapMan BIN categories was performed using Mercator version 3.6 (Lohse et al., 2014) together with MapMan BIN definitions as implemented in MapMan (Thimm et al., 2004). Raw count filtering and background definition were conducted in R (v4.3.2) using edgeR (v3.42.4). All expressed genes from the unfiltered count matrix were imported into edgeR, and lowly expressed genes were removed with the filterByExpr function, which adaptively selects expression thresholds while accounting for differences in library size. Genes retained after filtering were taken as the background universe and exported as gene IDs. Protein sequences for *Arabidopsis thaliana* (Araport11, June 2016 release) were downloaded from TAIR, and the peptide FASTA file was validated using the Mercator FASTA validator. Functional BIN assignments were generated with Mercator v3.6, and the resulting annotation table was cleaned in R to produce the final annotation reference. BIN enrichment for up-regulated and down-regulated gene sets was assessed using one-sided Fisher’s exact tests (performed in R) with the same filtered background gene universe supplied for both annotation and statistical testing. Because the Mercator v3.6 web interface could not complete the Fisher test step due to memory limitations, the enrichment was performed locally in R using identical background and annotation inputs to ensure reproducibility.

### In-gel NADP-ME activity assay

Approximately 100 mg of imbibed seeds from WT and *me1* lines was ground separately in liquid nitrogen and homogenized in 300 µl extraction buffer (100 mM Tris-HCl, pH 8.0; 100 mM NaCl; 0.5% (v/v) Triton X-100; 2 mM PMSF; 1% (w/v) PVP40). Samples were vortexed and centrifuged at 20,000 x g for 15 min at 4 °C, and the supernatant was collected in a clean tube. Protein concentration was determined using a modified Amido Black 10B precipitation method (Schaffner and Weissmann, 1973) with bovine serum albumin as standard for the calibration curve (Hüdig et al., 2022).

For native polyacrylamide gel electrophoresis (PAGE), 30 µg of protein per sample was loaded onto a non-denaturing 7% (w/v) polyacrylamide gel and electrophoresed at 100 V at 4 °C. Protein bands were visualized by an in-gel NADP-ME activity assay. Gel was first incubated in 50 mM Tris-HCl (pH 8.0) for 15 min at room temperature, followed by incubation in assay solution containing 50 mM Tris-HCl (pH 8.0), 10 mM L-malate, 10 mM MgCl_2_, 0.5 mM NADP, 0.05% (w/v) NBT, and 150 µM phenazine methosulfate (PMS). After a brief rinse with distilled water, gel was imaged under a scanner (CanoScan LiDE 400).

## Data analysis

All statistical analyses and data visualization were performed in R (version 4.4.1) using RStudio. Data were analyzed using linear models with heteroscedasticity-robust standard errors to account for unequal variances. Depending on the experimental design, one-way or two-way analysis of variance (ANOVA) was used to test for significant effects. When significant effects were detected, pairwise comparisons were performed using estimated marginal means with Sidak adjustment for multiple comparisons. Data visualization was generated using the ggplot2 package. For Figures 2B, 4, 5B-E, 6B, and Supplementary Figure 6, data were analyzed using two-way ANOVA testing the effects of line and treatment (or day where applicable). When significant effects were detected, pairwise comparisons among lines within treatments (or days) and among treatments within each line were performed using estimated marginal means with Sidak adjustment. For Supplementary Figures 1C, 2A, 8A, and 11, data were analyzed using one-way ANOVA followed by pairwise comparisons using estimated marginal means with Sidak adjustment. For Figure 9B, statistical differences between groups were assessed using the Watson-Williams test for equality of mean directions (Watson and Williams, 1956).

## Data availability

The data supporting the findings of this study are available from the corresponding author upon reasonable request. RNA sequencing data generated in this study have been deposited in the NCBI Sequence Read Archive (SRA) under BioProject accession PRJNA1418613 and are publicly available at https://www.ncbi.nlm.nih.gov/sra. The raw RNA-seq data (FASTQ files) associated with this study are accessible through this repository.

## Supplementary Data

**Supplementary Data 1. Differentially expressed genes identified in RNA-seq comparisons.** Excel file containing four worksheets listing significantly differentially expressed genes (adjusted *p* value < 0.05 and |log_2_FC| > 1) for the following comparisons: WT_ABA_vs_ctr, me1_ABA_vs_ctr, me1_vs_WT_ctr, and me1_vs_WT_ABA. Columns include Gene_ID (Arabidopsis locus identifier), log_2_FC, and Padj. Control (ctr) corresponds to ½ MS medium.

Supplementary Data 2. Functional annotation and MapMan classification of differentially expressed genes in *me1* compared with WT. Excel file containing worksheets listing genes uniquely up-or downregulated in me1 relative to WT under control (½ MS) and ABA conditions, together with their MapMan functional annotation. Columns include Comparison, Gene_ID, Gene name, log_2_FC, FC, Padj, MapMan bin, and Raw bin name. Additional sheets include intersecting gene sets and a summary of genes lacking MapMan bin assignments.

**Supplementary Data 3. Proteins associated with NADP-ME1 identified by co-immunoprecipitation and mass spectrometry.** Excel file containing proteins detected after co-immunoprecipitation (Co-IP) of NADP-ME1 followed by liquid chromatography–tandem mass spectrometry (LC-MS/MS). The file includes two worksheets corresponding to independent biological replicates (Replicate 1 and Replicate 2) and lists protein identifications together with associated mass spectrometry parameters, including peptide spectrum matches, intensity, log2 intensity, and differences of log2 intensities.

**Video 1.** Time-lapse sequence showing root growth of WT (right) and *me1.2* (left) seedlings imaged every 6 h after sowing on ½ MS plates. At 4 days after sowing, the lower part of the medium was replaced with ½ MS containing 200 mM NaCl, as indicated in the bottom right corner of each image.

## ACKNOWLEDGEMENTS

This work was supported by grants of the Deutsche Forschungsgemeinschaft to V.G.M. (MA2379/20-1) and by the University of Bonn through an Argelander Starter-Kit Grant to M.B. Confocal imaging was performed using a Leica TCS SP8 microscope at the IZMB funded by the DFG (INST 217/939-1 FUG). We gratefully acknowledge Dr. Fatima Chigri and Dr. Susann Frank for their technical support and training in the use of the Leica SP8 confocal microscope.

The authors have no competing interests (financial/non-financial) that might be perceived to influence the interpretation of the article.

## AUTHOR CONTRIBUTIONS

V.G.M. conceived the study. Y.F., M.K., E-M.S.I., M.M.S. G.P., V.W and M.S performed experiments and collected data. M.B., Y.F., S.M., and M.S. analyzed the data. V.G.M., M.B., and M.S. interpreted results. V.G.M, M.B., and Y.F. wrote the manuscript with input from all authors. V.M. supervised the project. V.G.M. and M.B. acquired funding.

## Notes

https://www.ncbi.nlm.nih.gov/sra

### Competing Interest Statement

The authors have declared no competing interest.

### Summary of Updates

We added to the acknowledgements: Confocal imaging was performed using a Leica TCS SP8 microscope at the IZMB funded by the DFG (INST 217/939-1 FUG). We gratefully acknowledge Dr. Fatima Chigri and Dr. Susann Frank for their technical support and training in the use of the Leica SP8 confocal microscope.

