## Supplementary Figures for "NADP-malic enzyme 1 couples ABA signaling to ROS-auxin patterning to restrict Arabidopsis root growth"

**Supplementary Figures Fu et al.**

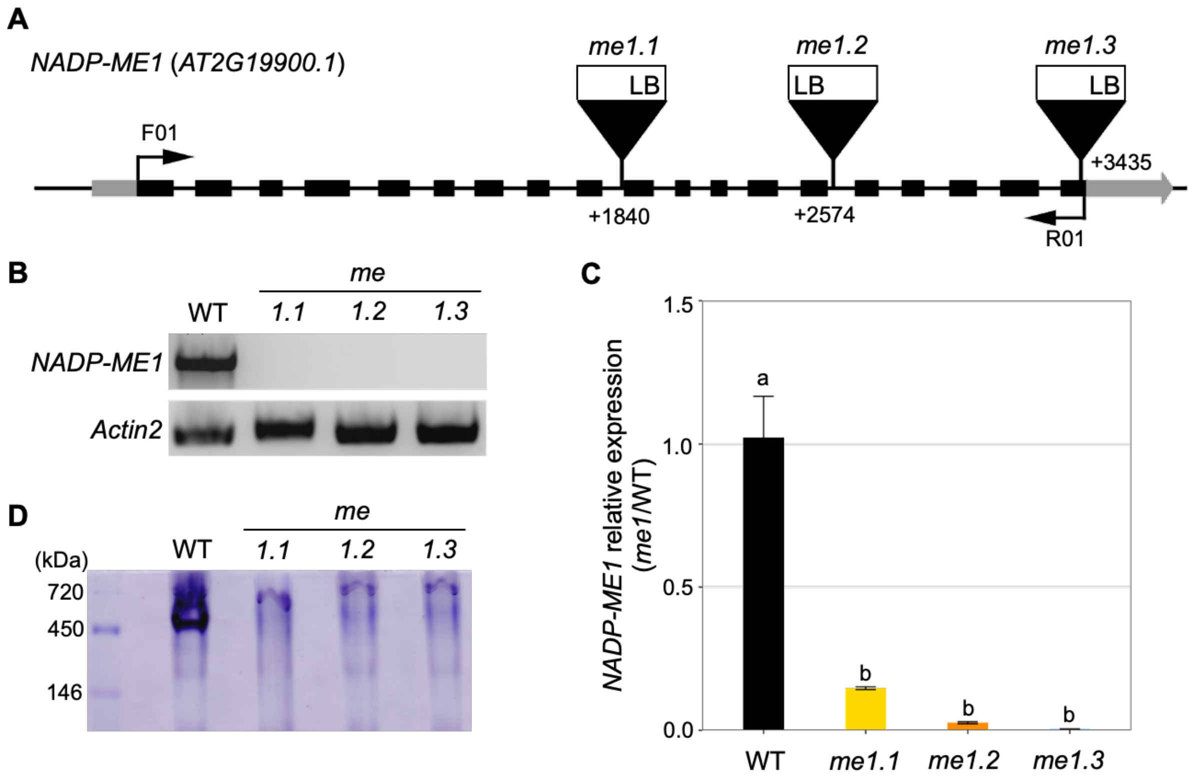


**Supplementary Figure 1. Characterization of *Arabidopsis NADP-ME1* T-DNA insertion lines. A)** Schematic representation of the *NADP-ME1* gene locus showing verified T-DNA insertion sites in three independent mutant lines: *me1.1* (SALKseq_047029.1), *me1.2* (SALKseq_036898.0), and *me1.3* (SALKseq_137251.1). Exons are depicted as boxes, with coding regions shown in black and untranslated regions in grey. **B)** RT-PCR analysis of full-length *NADP-ME1* transcripts in roots using primer pair F01 and R01. *Actin2* (AT3G18780) was used as an internal control. **C)** Relative *NADP-ME1* expression in roots of WT and the T-DNA insertion mutants determined by quantitative real-time PCR. Transcript levels were normalized to *Actin2*. Data represent means ± SE (n = 3). Differences among lines were assessed using one-way ANOVA. Different lowercase letters indicate significant differences among lines ($p < 0.05$). **D)** In-gel NADP-ME activity assay using extracts from seeds two days after imbibition of WT and the *me1* mutants.

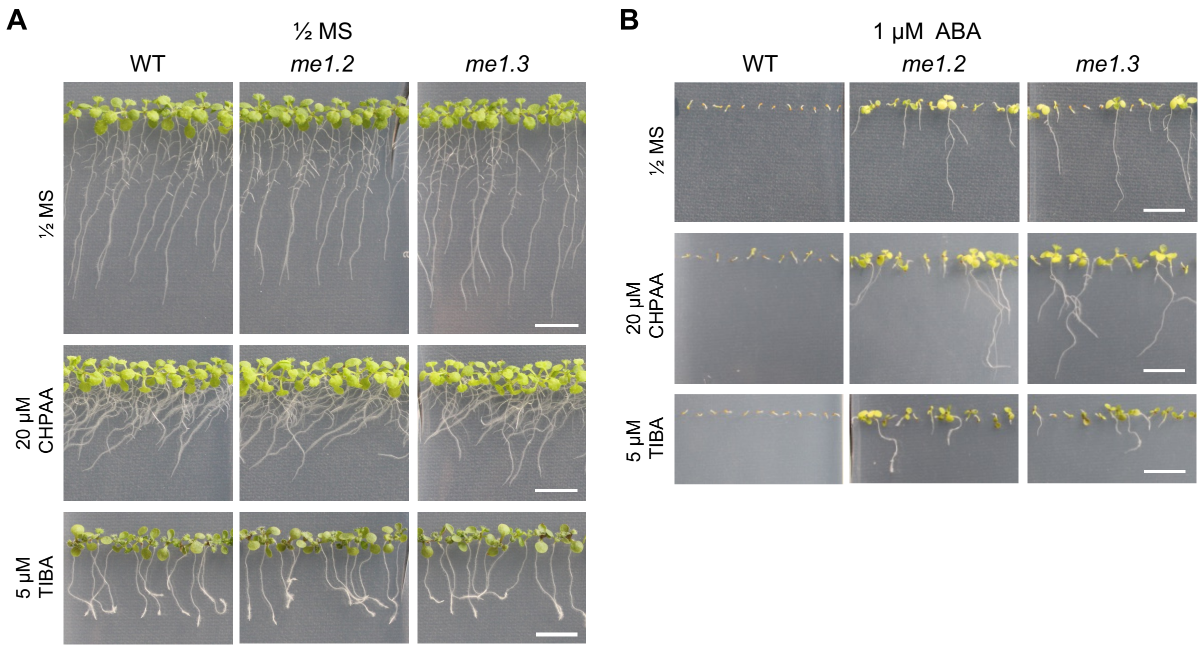


**Supplementary Figure 2. Auxin influx and efflux inhibitors altered *me1* root responses to ABA treatment. A)** Representative images of WT, *me1.2*, and *me1.3* seedlings grown for 10 days on ½ MS medium (control) or ½ MS supplemented with 20 $\mu$M CHPAA or 5$\mu$M TIBA. **B)** Representative images of WT, *me1.2*, and *me1.3* seedlings grown for 14 days on ½ MS medium containing 1 $\mu$M ABA alone or in combination with 20 $\mu$M CHPAA or 5 $\mu$M TIBA. Scale bar: 1 cm.

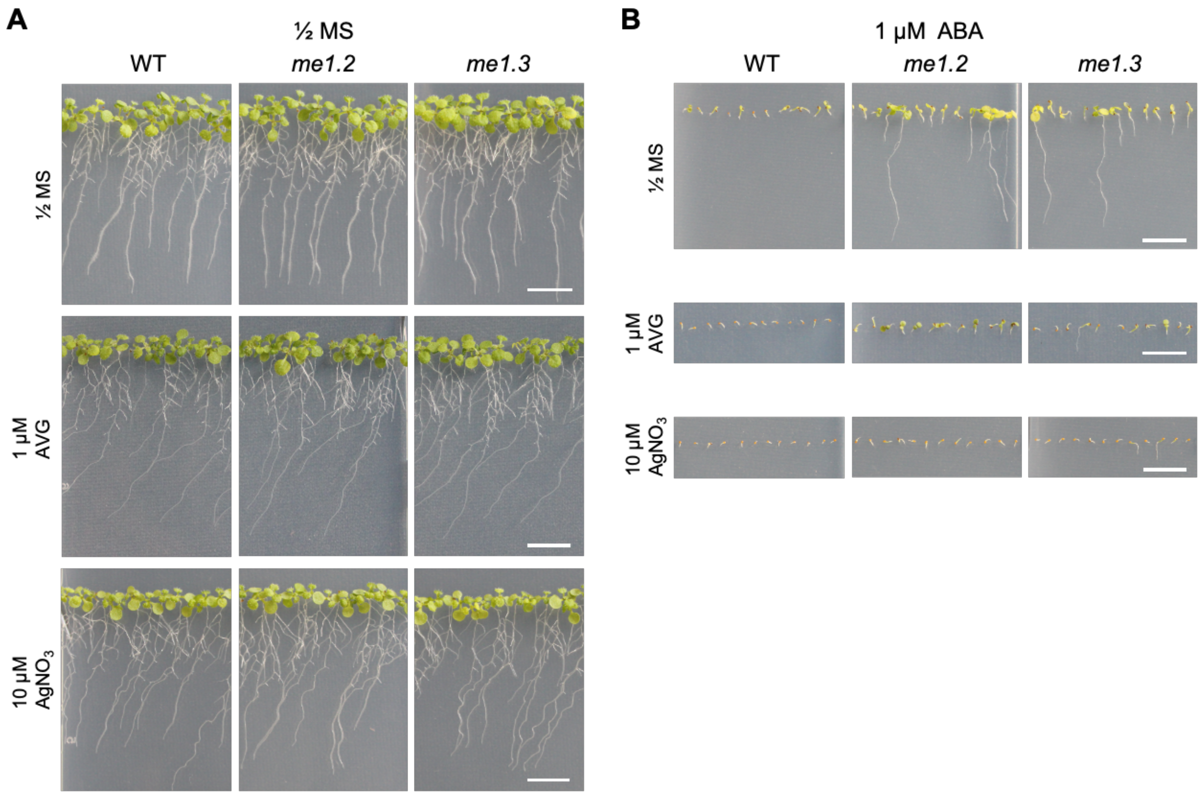


**Supplementary Figure 3. Ethylene biosynthesis and signaling inhibitors altered *me1* root responses to ABA treatment. A)** Representative images of WT, *me1.2*, and *me1.3* seedlings grown for 10 days on ½ MS medium (control) or ½ MS supplemented with 1 $\mu$M AVG, or 10 $\mu$M $AgNO_3$. **B)** Representative images of WT, *me1.2*, and *me1.3* seedlings grown for 14 days on ½ MS medium containing 1 $\mu$M ABA alone or in combination with 1 $\mu$M AVG, or 10 $\mu$M $AgNO_3$. Scale bar: 1 cm.

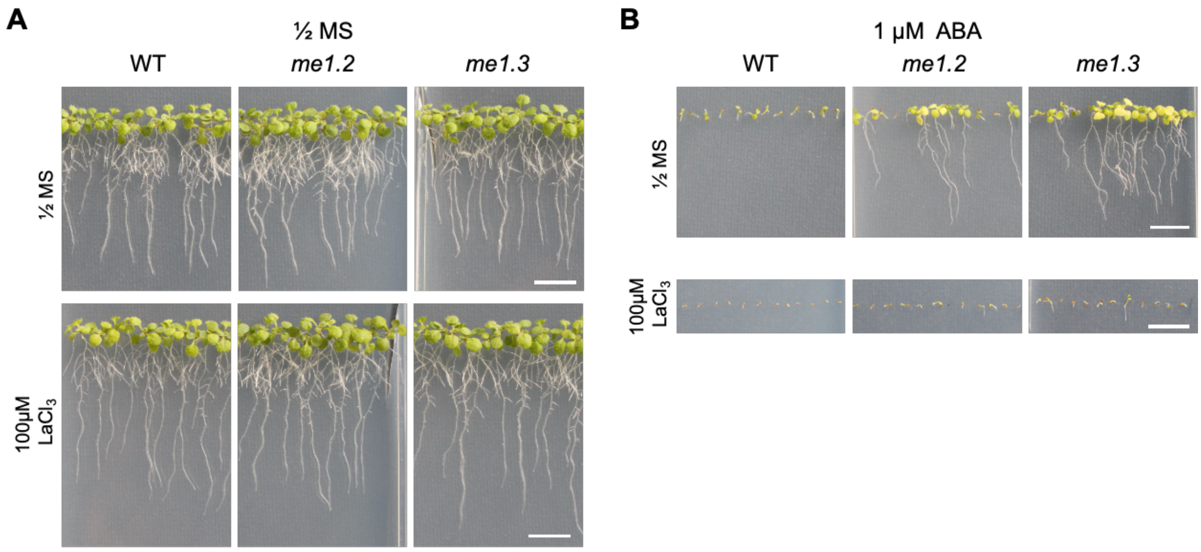


**Supplementary Figure 4. Calcium channel blocker altered *me1* root responses to ABA treatment. A)** Representative images of WT, *me1.2*, and *me1.3* seedlings grown for 10 days on ½ MS medium (control) or ½ MS supplemented with 100 $\mu$M $LaCl_3$. **B)** Representative images of WT, *me1.2*, and *me1.3* seedlings grown for 14 days on ½ MS medium containing 1 $\mu$M ABA alone or in combination with 100 $\mu$M $LaCl_3$. Scale bar: 1 cm.

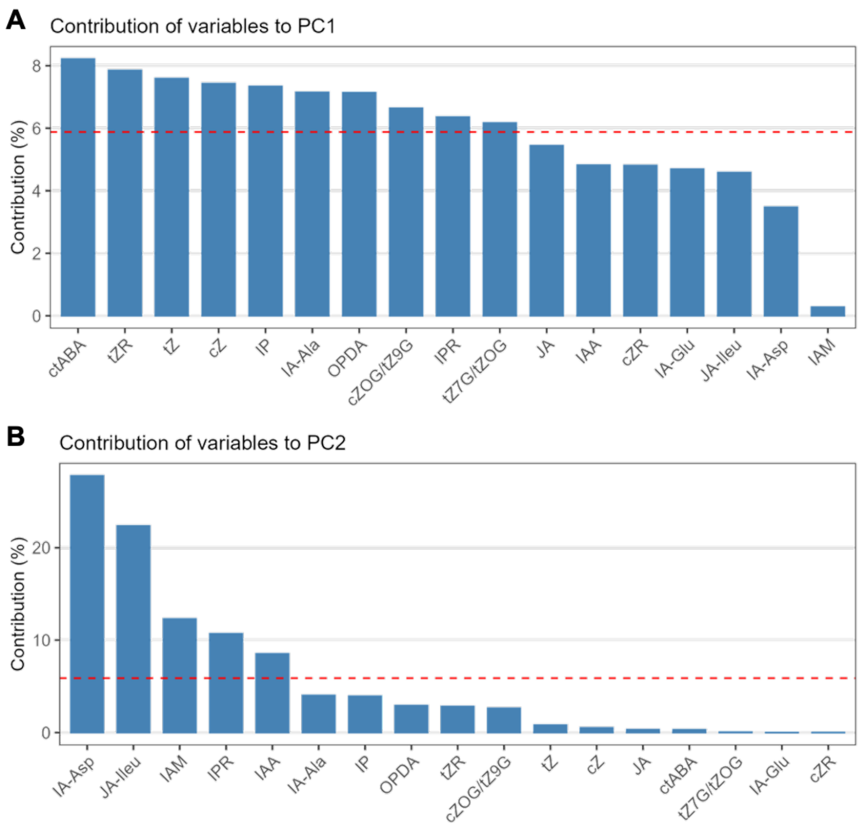


**Supplementary Figure 5**. Percentage contributions of phytohormones to PC1 and PC2 from the PCA shown in Figure 5. **A)** Contribution of individual phytohormones to the first principal component (PC1). **B)** Contribution of individual phytohormones to the second principal component (PC2). Bars indicate the percentage contribution of each phytohormone to the respective principal component.

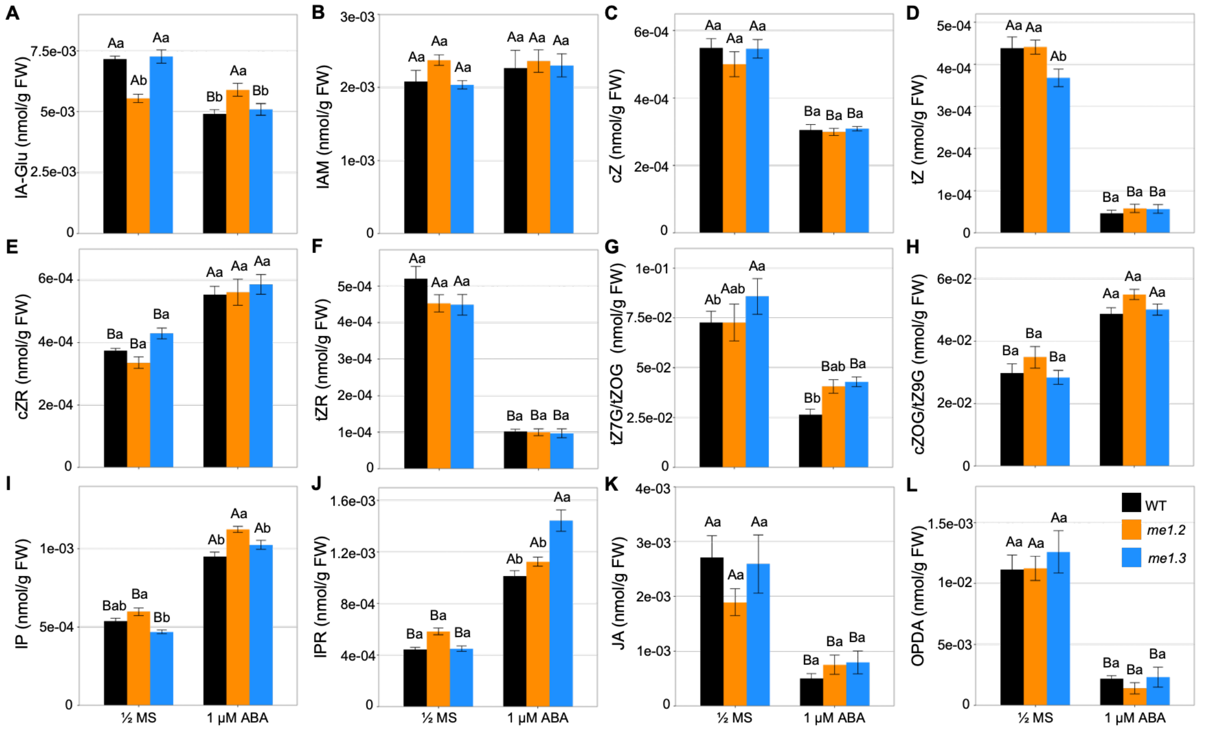


**Supplementary Figure 6. Phytohormone profiles of plants grown in the absence or presence of 1 µM ABA.** Seedlings of WT, *me1.2*, and *me1.3* were grown on ½ MS medium without or with 1 µM ABA for two days. Phytohormone concentrations were determined via targeted metabolite profiling. **A)** IA-Glu (indole-3-acetyl-glutamate); **B)** IAM (indole-3-acetamide); **C)** cZ (*cis*-zeatin); **D)** tZ (*trans*-zeatin); **E)** cZ-riboside; **F)** tZ-riboside; **G)** tZ7G/tZOG (*trans*-zeatin-7-glucoside/*trans*-zeatin-O-glucoside); **H)** cZOG/tZ9G (*cis*-zeatin-O-glucoside/*trans*-zeatin-9-glucoside); **I)** IP (isopentenyladenine); **J)** IPR (isopentenyladenosine); **K)** JA (jasmonic acid); **L)** OPDA (12-oxo-phytodienoic acid). tZ7G/tZOG and cZOG/tZ9G co-eluted and were not separated by HPLC. tZR and cZR, were quantified based on their peak areas relative to external calibration curves without the use of stable-isotope labelled standards. Data are presented as mean ± SE (n = 7). Differences were assessed using a two-way ANOVA (line and treatment), followed by pairwise comparisons among lines within each treatment and between treatments within each line using estimated marginal means with Sidak adjustment. Different lowercase letters indicate significant differences among lines within the same treatment, and different uppercase letters indicate significant differences among treatments within the same line ($p < 0.05$).

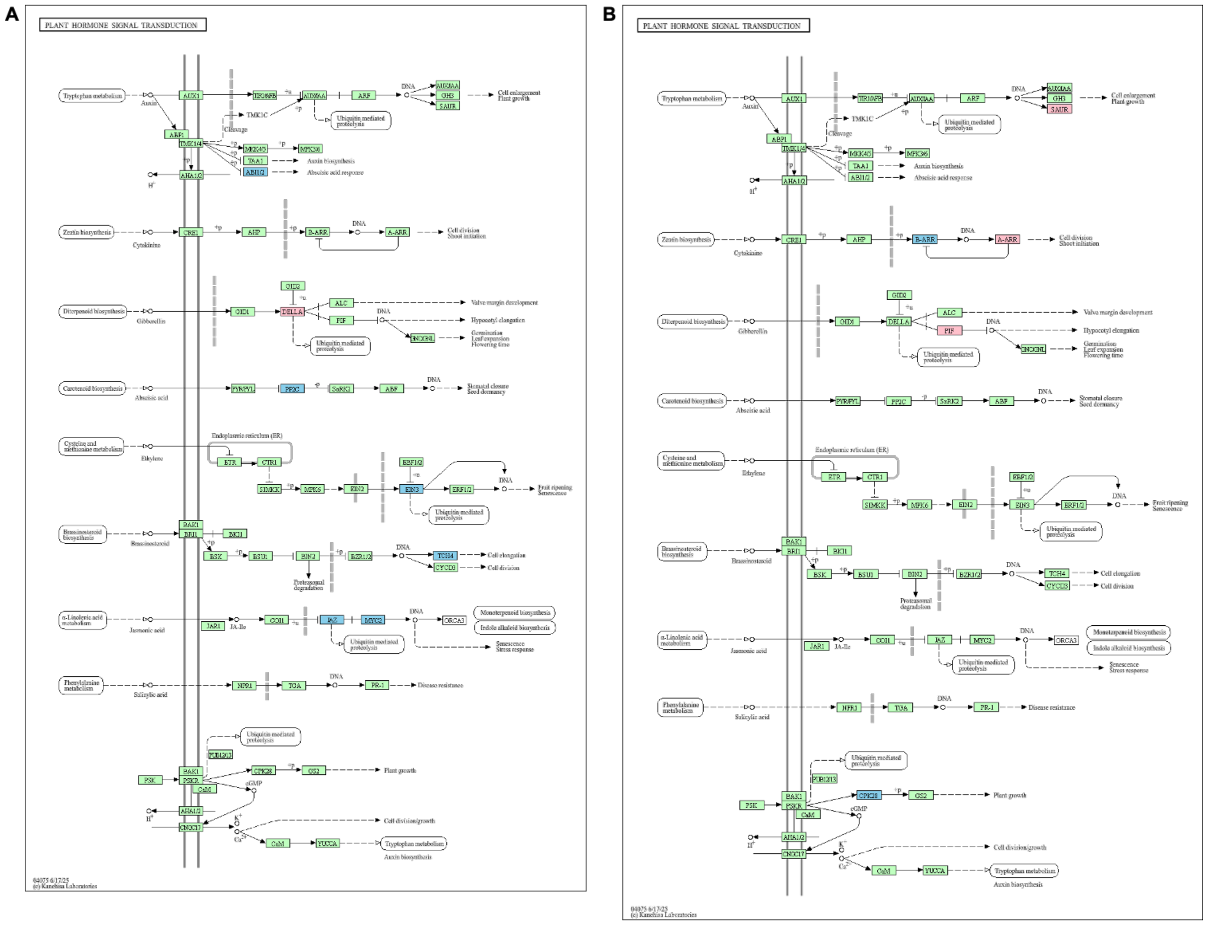


**Supplementary Fig. 7. Targeted KEGG pathway enrichment focusing on the “Plant hormone signal transduction” pathway in *nadp-me1* and WT under control and ABA conditions. A)** KEGG enrichment of the “Plant hormone signal transduction” pathway in the *nadp-me1* mutant compared with wild type under control condition. **B)** KEGG enrichment of the “Plant hormone signal transduction” pathway in *me1* compared with WT under ABA treatment. The diagram highlights up- and down-regulated genes (in red and blue, respectively) mapped to key hormonal signaling branches.

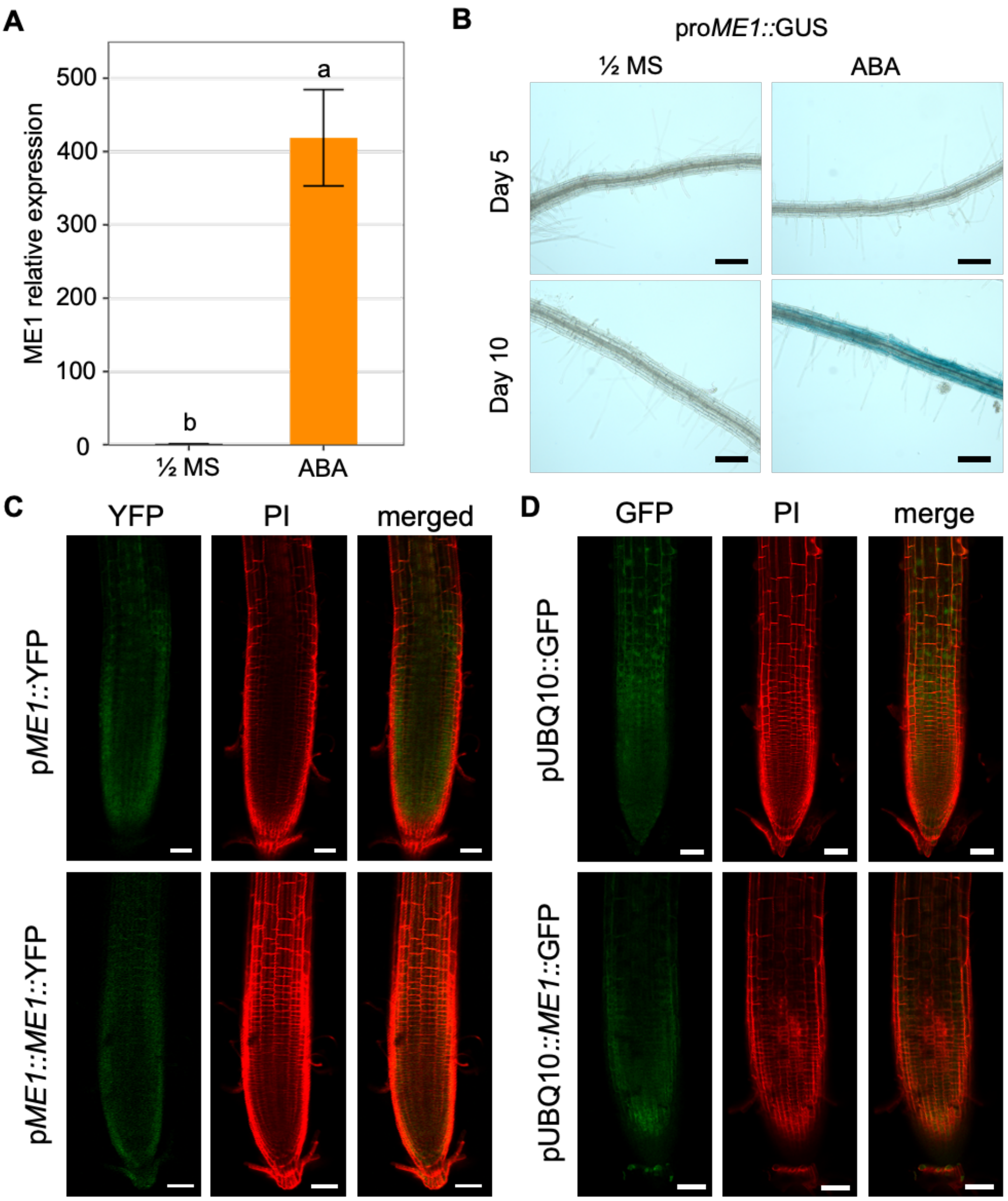


**Supplementary Figure 8. Expression analysis of *NADP-ME1* expression in roots under ABA treatment. A)** Transcript levels of *NADP-ME1* with or without 10 μM ABA treatment. Seedlings were grown on ½ MS medium for 10 days, then transferred to ½ MS medium with or without 10 μM ABA for an additional 12 hours. Data are presented as means ± SE (n = 4). Differences between treatments (½ MS medium with or without 10 μM ABA) were assessed using one-way ANOVA. Different lowercase letters indicate significant differences between treatments in WT ($p < 0.05$). **B)** Analysis of pro*ME1*::GUS expression during ABA treatment. Seedlings were grown on ½ MS medium for 5 and 10 days, then transferred to ½ MS medium with or without 10 μM ABA for an additional 12 hours. Scale bar: 100 μm. **C)** Seedlings were grown on ½ MS medium for 10 days and then transferred to ½ MS medium supplemented with 10 μM ABA for 12 h. Representative confocal images of seedlings expressing p*ME1*::YFP or p*ME1*::*ME1*::YFP were shown. YFP (yellow fluorescence) signal indicates transgene expression. **D)** Seedlings were grown on ½ MS medium for 10 days and then transferred to ½ MS medium supplemented with 10 μM ABA for 12 h. Representative confocal images of seedlings expressing pUBQ10::GFP or pUBQ10::*ME1*::GFP were shown. GFP (green fluorescence) signal indicates transgene expression. PI, propidium iodide staining; Merge, overlay of fluorescence channels. Scale bar: 50 μm.

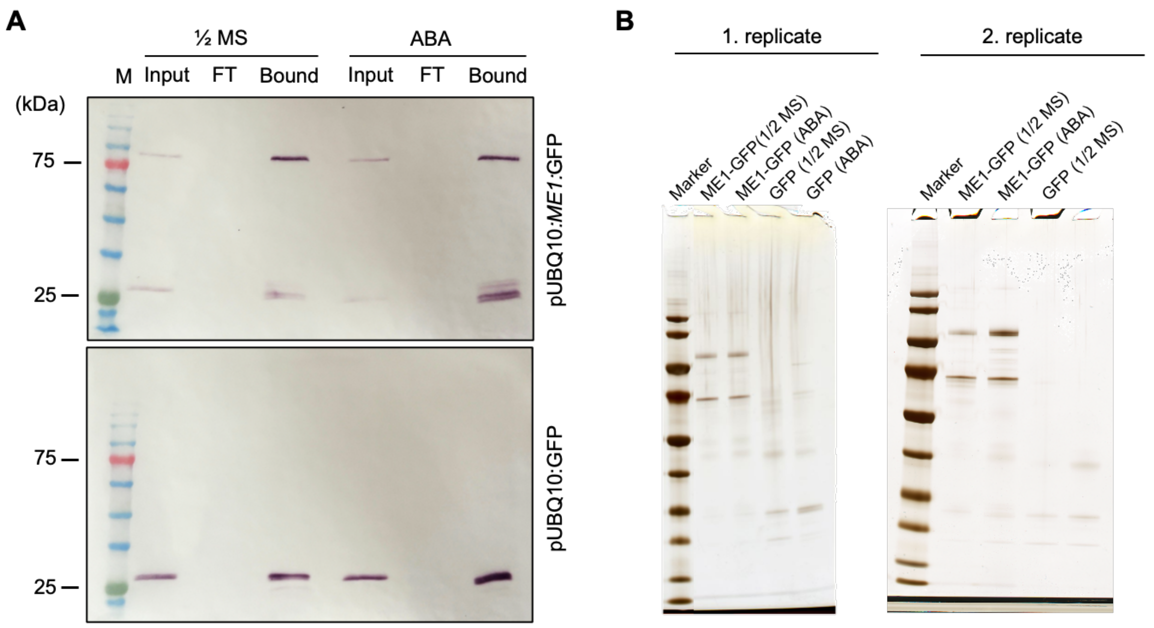


**Supplementary Figure 9. Co-immunoprecipitation analysis from roots under ABA treatment. A)** Protein fractions from co-immunoprecipitation (Co-IP) assays using root extracts from plants expressing pUBQ10::*ME1*::GFP and pUBQ10*::GFP*, grown on ½ MS medium for 10 days and then transferred to ½ MS medium with or without 10 $\mu$M ABA for an additional 12 hours. Proteins were immunoprecipitated using a mouse anti-GFP (IgG1κ) primary antibody and a goat anti-mouse IgG secondary antibody. M, ROTI®Mark Tricolor protein marker; FT, flow-through fraction. **B)** SDS-PAGE analysis of proteins co-immunoprecipitated with ME1-GFP and GFP from the bound fractions shown in (A), under control (½ MS) and ABA-treated (10 $\mu$M ABA) conditions. One-tenth of the total volume of each bound fraction was loaded per lane. Results are shown for biological replicates 1 and 2.

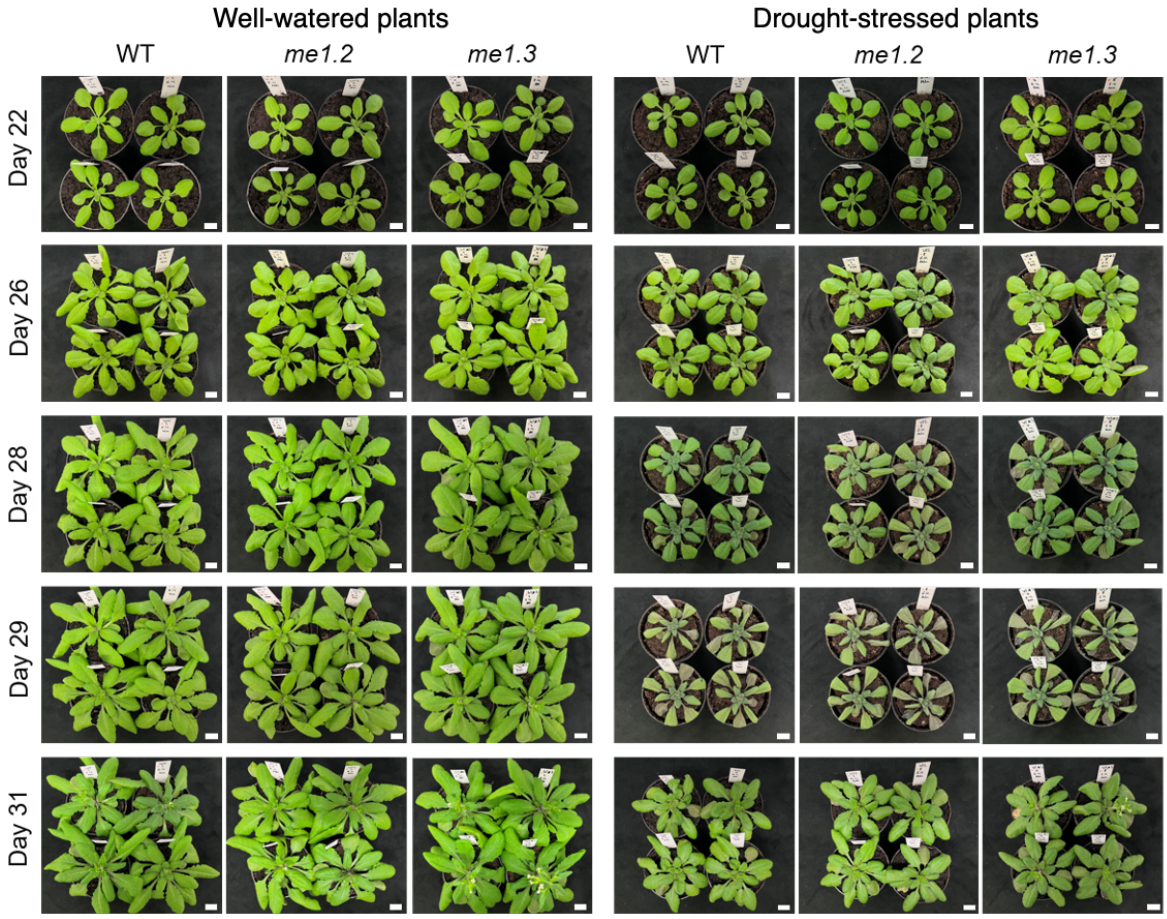


**Supplementary Figure 10. Phenotypes of WT and *me1* under well-watered and drought conditions.** Representative images of WT, *me1.2*, and *me1.3* seedlings on days 24, 26, 28, 29, and 31 are shown. Left panel: plants under well-watered conditions; right panel: plants under drought stress. For drought stress, plants were last watered on day 14 and re-watered on day 29 after imaging. Scale bars represent 1 cm.

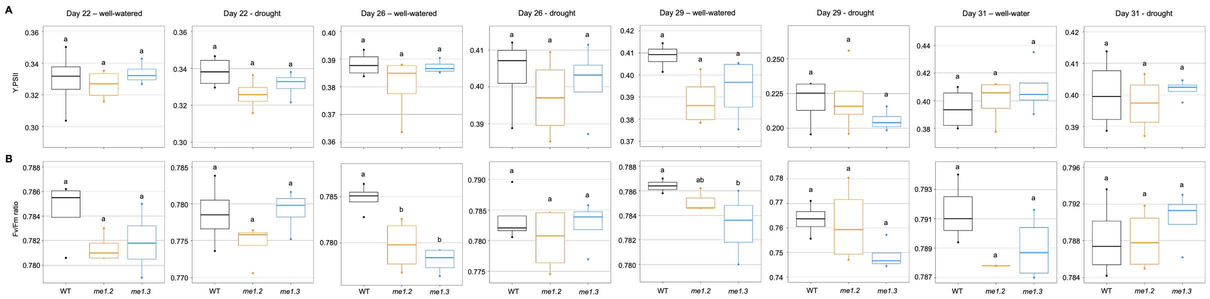


**Supplementary Figure 11. Time course analysis of PSII performance in WT and *me1* lines under control and drought conditions.** Boxplots show (**A**) the effective quantum yield of PSII (Y(II)) and (**B**) the maximum quantum efficiency of PSII (Fv/Fm) in WT, *me1.2*, and *me1.3* plants at different days under well-watered and drought stress. For each genotype, four biological replicates (plants) were analyzed, with five technical measurements per plant; mean values per plant were used for visualization. Differences among lines were assessed using one-way ANOVA. Different lowercase letters indicate significant differences among lines ($p < 0.05$).

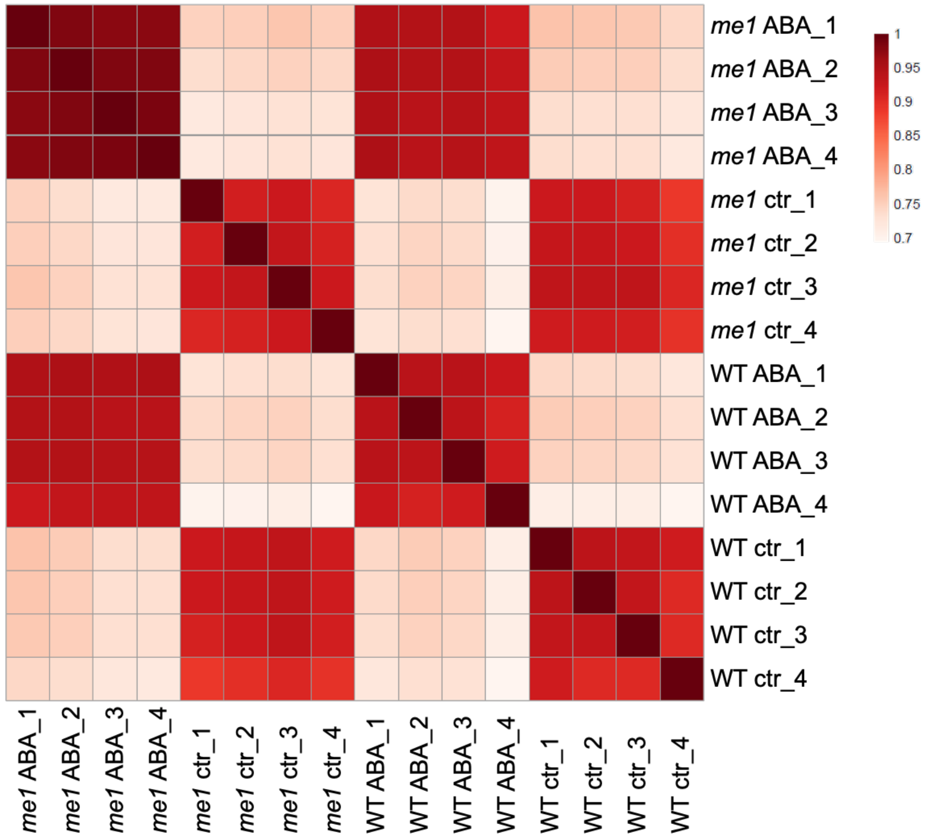


**Supplementary Figure 12. Spearman correlation heatmap** of RNA-seq samples based on **DESeq2 normalized read counts**, showing the similarity between biological replicates across genotypes (WT, *nadp-me1*) and treatments (control, ABA). Samples are displayed in a predefined order corresponding to genotype and treatment conditions. Control (ctr) corresponds to ½ MS medium.
